# Multi-objective domain adaptation for pan-cancer transfer of pharmacogenomic drug response to patient tumours

**DOI:** 10.64898/2026.11.04.764164

**Authors:** Teemu Rintala, Vittorio Fortino

## Abstract

Large-scale pharmacogenomic screens provide rich drug-sensitivity measurements in cancer cell lines, but transferring these measurements to patient tumours remains limited by biological heterogeneity and cross-domain distribution shift. Existing machine learning approaches, including adversarial autoencoder and domain-adaptation, address this gap by learning shared latent representations between preclinical models and patient samples. However, alignment without explicit biological constraints can produce unreliable representations when cancer-lineage structure and clinically relevant patient heterogeneity are not preserved. Here, we introduce MODAE+, a pan-cancer multi-objective deep learning domain-adaptation framework that jointly balances domain alignment, deconfounding, drug-response prediction, survival prediction, and cancer-lineage classification across 9,696 TCGA tumours, 1,293 CCLE cancer cell lines, and CTRP sensitivity profiles for 544 compounds in 738 matched cell lines. This multi-objective design improved cross-domain alignment while preserving cancer-lineage organization and patient heterogeneity. For drugs shared between the pharmacogenomic screen and TCGA treatment records, MODAE+ predicted sensitivity was significantly associated with overall survival among patients who received each drug in 46.3% of evaluated cancer–drug pairs by log-rank testing and 41.5% by multivariable Cox analysis. Biological validation across four curated drug–gene resources identified significant expression differences exceeding 1.5-fold for 34.9–64.4% of drugs in pan-cancer analyses, based on the lowest-p-value annotated gene per drug. Drug-class-specific patterns involved intended molecular targets for targeted therapies and predominantly transport, metabolism, detoxification and resistance-related genes for classical chemotherapies. As an additional validation, we derived pretreatment responder–nonresponder gene-expression signatures from 13 external breast cancer datasets and evaluated their concordance with MODAE+ predictions across 69 signature–drug comparisons in TCGA patients treated with paclitaxel or doxorubicin. Consensus signature scores correlated with predicted sensitivity at Spearman’s ρ = 0.791 and 0.724, respectively (mean = 0.758). Together, these complementary analyses provide supporting evidence for the clinical relevance and biological consistency of MODAE+ predictions, suggesting their potential to identify drug-sensitive tumour subgroups and generate hypotheses about mechanisms of drug resistance.

## Introduction

Predicting how individual patients will respond to anticancer therapies remains a fundamental challenge in precision oncology. In this context, large-scale pharmacogenomic screens in cancer cell lines, such as CCLE (Barretina *et al.*, 2012; Ghandi *et al.*, 2019), GDSC(Yang *et al.*, 2013; Iorio *et al.*, 2016), and CTRP (Seashore Ludlow *et al.*, 2015), have generated extensive drug-sensitivity measurements across hundreds of compounds, providing systematic in vitro resources for studying anticancer drug sensitivity and resistance. These datasets offer a unique opportunity to characterize therapeutic vulnerabilities in a controlled setting. However, bridging in vitro models with patient tumours, by leveraging transcriptomic profiles across bulk, single-cell, and, more recently, spatial modalities, remains a major obstacle to translating drug sensitivity predictions to real patients, as biological and technical differences between systems limit the reliability of inferred drug-response information at the patient level.

Early efforts to relate cell-line and patient transcriptomic data include linear domain adaptation methods, such as PRECISE (Mourragui *et al.*, 2019), which project both datasets into a shared low-dimensional subspace. Subsequently, batch-effect correction approaches, including ComBat and related strategies (Leek *et al.*, 2012), were applied to align cell-line and patient datasets (Salvadores *et al.*, 2020). More recent frameworks, such as Celligner (Warren *et al.*, 2021), perform alignment in a reduced-dimensional embedding space by combining contrastive principal component analysis with mutual nearest neighbor matching, thereby leveraging local transcriptional similarity rather than global statistical correction to reduce systematic differences between domains. These alignment strategies are also employed in downstream predictive tools, such as CellHit (Carli *et al.*, 2025), which integrates a modified Celligner pipeline prior to drug-response modelling. Despite reducing global dataset discrepancies, these approaches do not explicitly constrain alignment according to biologically corresponding cancer lineages across domains. As a result, cell lines may remain clustered in compact domain-specific regions rather than being anchored to patient tumours of matching biological origin, limiting cross-domain co-localization and hindering the transfer of drug-response information. Moreover, even when partial mixing is observed, the resulting embeddings may fail to preserve the molecular heterogeneity needed to identify clinically relevant patient subgroups with shared molecular patterns and consistent drug-response profiles. Deep learning approaches that do not rely on domain adaptation, through batch effect removal techniques, have also been proposed for drug sensitivity prediction. Examples include multi-omics representation learning frameworks such as MOLI (Sharifi Noghabi *et al.*, 2019), interpretable drug sensitivity models guided by deep learning, and graph-based architectures such as XGraphCDS (Wang *et al.*, 2024) and DeepCCDS (Wu *et al.*, 2025), which emphasize predictive performance, multi-omics integration, chemical structure modelling, or interpretability. However, these methods are typically trained within a single domain and do not explicitly address the discrepancy between preclinical models and patient tumours, limiting their ability to support cross-domain transfer.

To overcome this limitation, adversarial representation learning has emerged as a powerful paradigm for domain adaptation in pharmacogenomics. Methods such as AITL (Sharifi□Noghabi *et al.*, 2020) and CODE-AE-ADV (He *et al.*, 2022) employ adversarial training to reduce domain-specific discrepancies between cell-line and patient representations, thereby enabling transfer of drug-response information across domains. Related domain-adaptation strategies have also been extended to patient-derived xenograft models, as in TRANSPIRE-DRP (Yang *et al.*, 2025), which translates PDX-derived drug-response information to clinical patients using deep representation learning and adversarial adaptation. In particular, CODE-AE-ADV learns both shared and private latent components, allowing common molecular features to support transfer while retaining domain-specific variation separately. This is important because patient tumours and cell lines are not expected to become fully interchangeable in latent space, especially in pan-cancer settings where patient samples contain broader tissue- and tumour-specific structure. However, these approaches remain primarily optimized for drug-response transfer and do not explicitly constrain the shared latent space to preserve broader biological or clinical organization. Consequently, strong adversarial deconfounding may improve domain mixing while also weakening cancer-lineage structure and reducing the patient-level heterogeneity needed for clinically meaningful stratification.

Building upon this paradigm our previous work introduced MODAE, a multi-task deconfounding autoencoder that integrates cell-line and patient transcriptomic data while using survival prediction to preserve clinically informative patient variation (Rintala *et al.*, 2024). However, MODAE was validated in breast cancer using TCGA and SCAN-B data (Brueffer *et al.*, 2018), and did not explicitly address the pan-cancer problem of aligning cell lines with patient tumours of matching cancer lineage while preserving tumour-of-origin structure. This aspect remains insufficiently addressed by existing pharmacogenomic domain-adaptation approaches, which are primarily optimized for domain alignment and drug-response transfer. Extending the framework with multiple supervised objectives also creates an optimization challenge, as fixed loss weights may allow individual objectives to dominate training.

To address these challenges, we propose MODAE+, a pan-cancer extension of MODAE that incorporates explicit cancer-lineage supervision, an enhanced drug-response prediction architecture, and Fast Adaptive Multitask Optimization (FAMO) by Liu *et al*. (2023) to dynamically balance competing learning objectives during training. Together, these extensions are designed to align cell lines and patient tumours according to shared cancer-lineage structure while preserving biologically meaningful tumour organization and clinically relevant patient heterogeneity.

MODAE+ is trained on transcriptomic data from TCGA patient tumours (∼9,696 samples) and cancer cell lines from the CCLE (∼1,293 models), together with large-scale pharmacogenomic drug sensitivity profiles from CTRP, comprising 544 compounds screened across 738 cell lines (∼321,567 drug–cell-line pairs). The model learns to predict multi-drug sensitivity from gene expression while jointly optimizing survival and tissue classification objectives to constrain the latent space. Model performance is systematically evaluated using a combination of metrics assessing domain alignment, preservation of cancer-lineage structure, drug sensitivity prediction accuracy, and survival prediction. The clinical and biological relevance of the learned representations and predictions is then further assessed through downstream validation analyses. In previous studies, these have relied largely on binary drug responder classification accuracy (He *et al.*, 2022; Yang *et al.*, 2025). However, responder labels are challenging to define, and observer variability remains despite standardization (Smesseim *et al.*, 2026). We present and implement a more robust approach using treatment information, patient survival, and predicted drug sensitivities directly by testing whether predicted drug sensitivities are predictive of patient survival and whether they can stratify patients into groups with distinct outcomes. Additionally, we show that predicted sensitivity is associated with differential expression of corresponding drug target genes. Finally, using gene-expression signatures derived from pretreatment responders versus nonresponders in external breast cancer cohorts we assessed whether MODAE+ predictions agree with clinically observed response-associated expression patterns.

## Materials and methods

### Training datasets and preprocessing

MODAE+ was trained using gene-expression profiles from human tumours and cancer cell lines, experimental drug-sensitivity measurements from cell lines, and patient survival and cancer-lineage annotations. TCGA provided the patient data, CCLE the cell-line expression profiles, and CTRP the drug-response measurements, enabling the model to learn transferable representations and estimate drug sensitivity in patient tumours. Pre-processed TCGA patient RNA-Seq data were obtained via CuratedTCGAData [v2.0.1] (Ramos *et al.*, 2020), restricted to primary solid tumours, and pre-processed CCLE cell-line RNA-Seq data from the DepMap portal(Corsello *et al.*, 2020), restricted to solid lineages. TCGA profiles were log2(TPM+1)-transformed (DepMap profiles were already log-scaled); genes with average expression below 1.0 in either dataset were removed, and patient and cell-line profiles were independently standardized to zero mean and unit variance, yielding 9,696 patient profiles, 1,293 cell-line profiles, and 12,214 genes. Curated survival data were obtained from Liu *et al*. (2018). Unlike our previous work (Rintala *et al.*, 2024), we used reprocessed CTRP drug-sensitivity data from Xia *et al*. (2022) who refit response curves and computed drug-sensitivity scores across sources; this comprised 544 drugs screened across 738 expression-matched cell lines (321,567 drug–cell-line pairs). Because the model predicts sensitivity from gene expression alone, and each drug has a corresponding output unit, cross-validation was performed across cell lines rather than drugs, with missing values masked from all loss and evaluation terms. Furthermore, the CV folds were stratified with respect to the presence of any drug sensitivity data on cell-lines and 10 survival time quantiles on patients to evenly distribute the data.

### MODAE-plus: an adaptive multi-objective deep latent spaces for tissue-aware drug response prediction

MODAE+ is a multi-objective deep-learning framework that jointly models cancer cell lines and patient tumours from gene expression within a shared latent representation, preserving molecular features common to both domains while transferring drug-response information learned from cell lines to patient tumours. It comprises a shared encoder, a decoder, a domain discriminator, and three supervised heads for drug sensitivity, patient survival, and cancer-lineage classification. The encoder projects both patient and cell-line profiles into a latent representation explicitly partitioned into shared and domain-specific (private) components, and reconstruction uses masked latent representations so that each domain is reconstructed only from the shared embedding and its own private embedding. To reduce domain-specific differences, the shared embeddings are aligned adversarially using the Wasserstein GAN framework (Arjovsky *et al.*, 2017; Gulrajani *et al.*, 2017), and a deconfounding norm penalty that enforces orthogonality between shared and private embeddings, isolating dataset-specific features into the private embeddings (He *et al.*, 2022). The shared representation then drives the three supervised tasks: drug sensitivity via a multi-channel head that learns drug-specific representations, patient survival via a Cox proportional-hazards objective, and an auxiliary cancer-lineage classification based on OncoTree Level 1 annotations [version 2021_11_02] (**Supplementary Tables S1–S2**) that preserves biologically meaningful organization across both domains. Unlike the original MODAE, which used manually tuned fixed loss weights, MODAE+ applies Fast Adaptive Multitask Optimization (FAMO; Liu *et al*. (2023)) to dynamically balance the competing objectives, including patient and cell-line reconstruction, adversarial alignment, deconfounding regularization, drug-sensitivity prediction, survival prediction, and cancer-lineage classification, thereby preventing any single objective from dominating. Reconstruction and lineage-classification losses are computed separately for patient and cell-line samples, allowing domain-specific control during optimization. The pseudocode of the MODAE+ algorithm is provided in **Supplementary Information File**.

### Validation framework and selection of the final latent space model

Model hyperparameters were optimized using a random search strategy in combination with nested cross-validation, with the ranges and candidate values explored for each hyperparameter reported in **Supplementary Table S3**. Each randomly sampled hyperparameter configuration was evaluated independently, with the inner cross-validation loop used for hyperparameter selection and the outer loop providing an unbiased estimate of model performance. The resulting performance metrics were used to rank the evaluated configurations and identify the optimal model for downstream analyses. A summary of the cross-validation results for all evaluated configurations is provided in **Supplementary Table S4**. The final hyperparameter configuration used for downstream analyses was selected according to a weighted optimization score combining the mean patient survival concordance index, the top 10th percentile drug-sensitivity, cancer lineage classification balanced accuracy, and the negative domain prediction balanced accuracy obtained across the cross-validation folds. This composite score was designed to favour models that simultaneously preserved clinically relevant patient representations, accurately transferred drug-response information, retained biologically meaningful cancer lineage structure, and minimized domain-specific information in the shared latent space. To avoid objective misalignment, the domain predictions for evaluation were performed with an independent random forest classifier.

### Optimization settings

Model training was performed using mini-batches of 32 samples and 5-fold cross-validation, with each model trained for a maximum of 150 epochs. The original module-wise pretraining strategy described in the original MODAE framework was disabled, and all network components were optimized jointly throughout training. Model optimization was performed using the Adam optimizer, while separate Adam optimizers were used for the adversarial domain critic, the deconfounding correction module, and the FAMO weights. For the model and adversarial optimizers, the momentum parameters were fixed toβ_1_ = 0.0 and β_2_ = 0.9 while learning rate was tuned with random search. Following FAMO guidelines, the weight optimizer was set to use β_1_ = 0.9, β_2_ = 0.999, and a learning rate of 0.025 while a weight decay parameter was applied and tuned with random search. Adversarial domain alignment was implemented using the Wasserstein loss with a gradient penalty of 1.0. Then, FAMO was used to dynamically balance the eight optimization objectives described above.

### Pharmacological class annotation

To support interpretation of the downstream validation analyses at the level of broader therapeutic mechanisms, drugs were grouped into pragmatic, mechanism-based pharmacological classes specifically for the purposes of this study. Classes were defined according to the predominant therapeutic mechanism or mode of action of each compound, grouping drugs with related pharmacological mechanisms into broader categories suitable for class-level comparison. This classification was subsequently used in both the treatment-specific survival analysis and the drug-target expression analysis, allowing results to be summarized beyond individual compounds and compared across broader modes of action. The complete drug-to-pharmacological-class mapping is reported in **Supplementary Table S5.**

### Drug sensitivity model transfer evaluation using survival data

To assess the clinical relevance of predicted drug sensitivities, we analysed survival in treated TCGA subsets, using treatment information from He *et al*. (2022) and updated TCGA survival annotations from Liu *et al*. (2018). Each cancer–drug pair was analysed separately, including patients who received the drug as a single agent or within a combination; survival times were calculated from the day of first treatment administration by adjusting the recorded survival time according to treatment initiation. Both overall survival (OS) and progression-free interval (PFI) were evaluated using three complementary strategies. First, predicted sensitivity was assessed as a continuous predictor via Harrell’s concordance index (C-index), testing whether patients predicted more sensitive experience more favourable outcomes: values above 0.5 indicate positive concordance, ∼0.5 no prognostic information, and below 0.5 an inverse association. Second, patients were stratified into predicted-sensitive and predicted-resistant groups (drug-specific score above or below the mean) and compared by Kaplan–Meier curves and log-rank tests. Third, predicted sensitivity was included as a continuous variable in multivariable Cox proportional-hazards models with age at diagnosis and AJCC stage as covariates, quantifying its independent contribution while accounting for clinical confounders (significance by Wald-test p-values); covariates with no variation within a cohort were excluded. Treatment information was used exclusively for validation, never during training. For comparison, the same framework was applied to CODE-AE-ADV using the authors’ class-probability predictions, with a threshold of 0.5 used to define sensitive and resistant groups. Because drug panels and cohorts differed between studies, main-text comparisons were restricted to overlapping cancer–drug pairs. The complete cancer–drug-specific survival-validation results are reported in **Supplementary Table S6**.

### Differential drug target gene expression analysis

To further assess the biological plausibility of the transferred drug-sensitivity predictions, we evaluated differential expression of drug-target genes between patients predicted sensitive versus resistant to a given treatment, reasoning that biologically meaningful predictions should associate with alterations in the drug’s target genes, a strategy also used by He *et al*. (2022) and in our earlier work (Rintala *et al.*, 2024). Drug targets were downloaded from the Comparative Toxicogenomics Database (Davis *et al.*, 2021), DrugBank (Wishart *et al.*, 2006, 2018; Knox *et al.*, 2024), PharmGKB (Gong *et al.*, 2021), and OpenTargets(Ochoa *et al.*, 2023). Patients were then stratified into predicted-sensitive and predicted-resistant groups according to whether their MODAE+ score for a given drug was above or below the drug-specific mean. Target-gene expression differences between the two groups were assessed using two-sided t-tests. For each drug–target gene–cancer association, we recorded the mean target-gene expression and number of samples in the predicted sensitive and resistant groups, the nominal p-value, the log□fold-change calculated as resistant versus sensitive, and the target-annotation source. Positive log□FC values therefore indicate higher target-gene expression in predicted resistant tumours, whereas negative values indicate higher expression in predicted sensitive tumours. The complete set of individual associations is reported in **Supplementary Table S7**. To summarize the extent of target-related support across drugs and cancers, these individual associations were subsequently evaluated using four complementary association-set strategies. In the *cancer-specific strategy*, all tested drug– target associations within individual cancer types were retained, whereas in the *pan-cancer strategy* all corresponding associations across all tumours were retained. Two additional *minimum-p-value strategies* were then used to assess whether at least one annotated target-associated gene supported the predicted sensitivity pattern. For *the minimum cancer-specific strategy*, the annotated target gene with the smallest nominal p-value was retained for each drug–cancer pair, separately within each target-annotation source. For *the minimum pan-cancer strategy*, instead, the annotated target gene with the smallest nominal p-value was retained for each drug, again separately within each target-annotation source. For each strategy and annotation source, the significance rate was calculated as the proportion of retained associations satisfying both nominal p < 0.05 and |log□FC| > log□(1.5).

### External breast cancer response signature concordance

As an additional post hoc validation, we evaluated whether MODAE+ drug-sensitivity predictions were concordant with treatment-response expression signatures derived from external breast cancer cohorts available through the Gene Expression Omnibus (GEO). The analysis included 13 GEO accessions comprising four RNA-seq and nine microarray datasets, organized into 40 treatment-, endpoint-or receptor-subtype-specific source groups. The primary analysis comprised 69 signature–drug comparisons: 34 for paclitaxel and 35 for doxorubicin (**Supplementary Tables S8–S11**). All eligible comparisons within this subset were reported, irrespective of their correlation with MODAE+ predictions or their source-cohort classification performance. The source collection comprised GSE123845, GSE164458, GSE194040, GSE20194, GSE20271, GSE207248, GSE22226, GSE23988, GSE25055, GSE25066, GSE319641, GSE32646 and GSE41998. GSE194040 (I-SPY2) contributed 22 treatment-arm–receptor-subtype groups. Each source group required at least three patients per outcome class, with one pretreatment specimen selected per patient. Source-group characteristics, sample numbers, treatment regimens and response definitions are provided in **Supplementary Table S8**. GEO accessions were not necessarily independent; GSE25055 and GSE25066 contain overlapping patient material. Expression matrices were analysed on dataset-appropriate log scales, applying log (TPM + 1) where TPM values were available and retaining previously log-normalized data without further transformation. Dataset-specific transformations, expression units and download information are documented in **Supplementary Table S9**. Then, clinical response groups were defined using cohort-provided outcome labels. For cohorts reporting pathological response, patients achieving pathological complete response (pCR) were classified as responders and those with residual disease as nonresponders. For cohorts using clinical response, complete or partial response (CR/PR) defined responders, with nonresponders assigned according to the cohort-specific criteria documented in **Supplementary Table S8**. Additionally, comparisons were labelled exploratory ([E]) when the available treatment metadata did not permit confident attribution of the source response signature to the individual drug being evaluated. This classification was based on treatment documentation, independently of correlation strength or statistical significance; cohort-specific explanations are provided in Supplementary Table S8. Differential expression was assessed between pretreatment samples from responders and nonresponders, using these cohort-specific response definitions, and restricted to the analysis to genes shared between each source dataset and TCGA. For each gene ***g***, an unmoderated equal-variance two-sample t-test produced a signed statistic (***t_g_***), with positive values indicating higher expression in responders. Genes were ranked by ***|t_g_|***, with ties resolved by gene name. The primary signature contained the 200 highest-ranked genes, with top-10 and top-50 signatures evaluated as sensitivity analyses. Gene weights and patient scores were calculated as: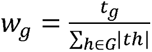 ; 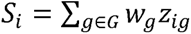 ; where (***G***) denotes the selected gene set and (***z_ig_***) denotes standardized expression of gene ***g*** in patient ***i***. Positive weights therefore represented genes more highly expressed in responders, whereas negative weights represented genes more highly expressed in nonresponders. Constant features were omitted during standardization. No statistical-significance threshold was imposed for gene selection, and score directions were not reversed after examining their correlations with MODAE+. Additionally, we also used pathway-restricted signatures based on the MSigDB Hallmark MITOTIC_SPINDLE (labelled as ‘Spindle DE’), DNA_REPAIR (labelled as ‘Repair DE’) and E2F_TARGETS (labelled as ‘E2F DE’) gene sets (version 2025), respectively to assess whether processes relevant to paclitaxel activity, doxorubicin-induced DNA damage and proliferation, respectively, strengthened concordance with MODAE+ predictions. Gene weights were then calculated from the same responder–nonresponder differential-expression statistics described above. The ability of each signature to distinguish responders from nonresponders within its source cohort was assessed using five repeats of stratified cross-validation (25 tests in total). In each fold, gene ranking and selection, gene weights, and expression means and standard deviations were estimated exclusively from training patients. For pathway signatures, gene-set membership was predefined, while weights and standardization parameters were estimated from the training patients. Each signature’s ability to distinguish responders from nonresponders within its external source cohort was evaluated using ROC AUC in held-out patients, averaged across cross-validation folds and repeats. Final gene selection and weights were then estimated using all patients in each source group, and the resulting signatures were applied to TCGA expression profiles to assess concordance with MODAE+ predictions. The signatures were applied to TCGA breast cancer subgroups comprising patients who had received paclitaxel or doxorubicin and had available expression profiles, MODAE+ predictions and treatment-start information. Drug–cancer groups required at least 30 recipients with non-zero predictions, after which zero-valued predictions were also retained. Eligible patients were split into calibration and evaluation subsets (60:40; seed 20261021), yielding 129/83 paclitaxel recipients and 186/130 doxorubicin recipients. Since signature scores depend on both the gene weights derived from external cohorts and standardized expression values, gene-specific means and standard deviations were estimated from the TCGA calibration subset and applied unchanged to evaluation patients. Scores were then calculated as weighted sums of standardized expression. Spearman correlations with original MODAE+ predictions were finally calculated for 34 paclitaxel and 35 doxorubicin top-200 signature comparisons, with top-10, top-50 and pathway-restricted signatures assessed additionally. Source-cohort AUCs and TCGA correlations are reported in **Supplementary Tables S10** and **S11**, respectively. To summarize agreement across external signatures, a consensus score was calculated for each TCGA patient and drug. First, individual signature scores were first standardized using means and standard deviations estimated from calibration patients. Subsequently, scores from source groups belonging to the same GEO accession were averaged, followed by averaging across accessions with equal weights. This prevented accessions contributing more source groups from dominating the consensus. Then, the resulting patient-level consensus scores were correlated with MODAE+ predictions in held-out evaluation patients using Spearman correlation. Overall concordance was summarized as the arithmetic mean of the two drug-specific consensus correlations, giving equal weight to each drug.

## Results

### MODAE+ resolves the trade-off between domain alignment and lineage preservation in cell-line–tumour integration

Because MODAE+ optimizes several competing objectives at once, no single hyperparameter setting is best for all of them, so we searched a large space of configurations and evaluated each across four axes: domain alignment (balanced accuracy of a random-forest batch classifier, RFC-batch BACC, where values near 0.5 mean cell lines and patients are indistinguishable), cancer-lineage preservation (CL-tissue BACC), drug-sensitivity prediction (R²), and survival prediction (C-index). The resulting Pareto fronts (**Supplementary Figure S1**) revealed distinct trade-off structure. Configurations that improved drug-sensitivity prediction generally also achieved higher survival performance, indicating that these two clinically oriented objectives are largely compatible and can be optimized together. Domain alignment, however, stood in tension with predictive performance: the strongest drug-response and survival models tended to retain some residual batch separability rather than driving it fully to chance, so that the best overall configurations occupy a compromise region of the Pareto front rather than the extreme of any single axis. Cancer-lineage preservation improved together with better alignment, but only partially tracked the predictive objectives, reflecting the intrinsic difficulty of flattening domain differences while retaining the biological structure that distinguishes tumour types.

Guided by these fronts, we selected the configuration that maximized the weighted training objective (marked with an asterisk in **Supplementary Figure S1**), which represents a balanced solution that combines strong drug-sensitivity and survival prediction with near-complete domain alignment (RFC-batch BACC ≈ 0.61), and used it for all downstream analyses. Its test-set performance (**Supplementary Figure S2**) demonstrates improved CCLE–TCGA alignment and competitive survival prediction, while the lower held-out drug-response R² is consistent with the known limited generalization of drug-response models on independent test data (Sinkala *et al.*, 2020).

We then compared the integrated space of the selected model against standard alternatives (**Figure 1A–E**), in which pan-cancer TCGA tumours (crosses) and CCLE cell lines (circles) are coloured by tissue of origin. In the unadjusted data (**Fig. 1A**), the two domains separated almost completely. ComBat correction (**Fig. 1B**) reduced these global differences but did not align cell lines with tumours of the same lineage. Celligner (**Fig. 1C**) co-localized some tissues but remained inconsistent across cancer types, and CODE-AE-ADV (**Fig. 1D**) removed the domain signal only by collapsing the samples into a continuous, regular structure with no recognizable tissue organization. MODAE+ (**Fig. 1E**) instead co-localized cell lines and patient tumours by cancer lineage, grouping samples of the same tissue together regardless of their domain of origin.

To quantify lineage recovery from each embedding, we then classified samples by tissue of origin using a k-nearest-neighbour classifier, for which higher accuracy indicates better-retained lineage structure (**Fig. 1F**). MODAE+ achieved the highest balanced accuracy (0.92) and per-lineage F1-scores of 0.70–0.99 across most tissues. CODE-AE-ADV ranked lowest, with balanced accuracy of 0.15 and F1 below 0.25 for most lineages (undefined for the rarest), quantifying the over-deconfounding evident in its UMAP. The ComBat variants gave intermediate lineage recovery while incompletely removing domain effects, and Celligner improved specific tissues but stayed inconsistent across cancer types. Ablations isolated the contribution of each MODAE+ component: removing cancer-lineage supervision (MODAE_NO_TIS_CL) or deconfounding (together with private encodings) (MODAE_NO_PE_NO_DEC) sharply reduced lineage recovery, and removing adaptive loss balancing (MODAE_NO_FAMO) destabilized performance across tissues. Across all methods and ablations, the full MODAE+ model gave the best combined alignment and lineage recovery, retaining tumour-of-origin structure while closing the domain gap.

**Figure 1.**
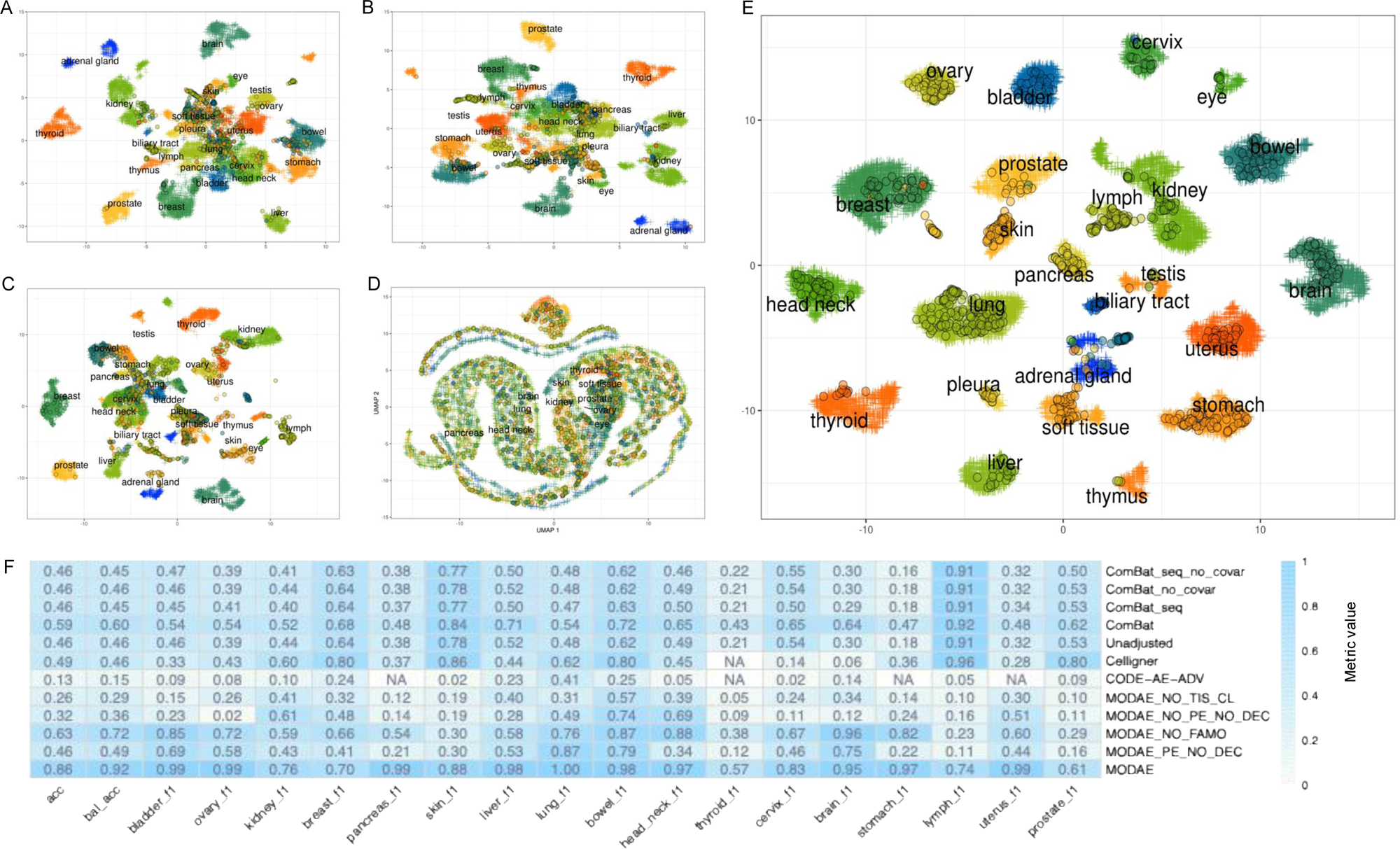
Comparison of alignment strategies for integrating cell-line and patient transcriptomic data. (A–E) UMAP projections of pan-cancer TCGA samples and CCLE cell lines under different integration strategies: (A) unadjusted bulk RNA-seq expression data, (B) ComBat-adjusted data, (C) Celligner, (D) CODE-AE-ADV and (E) MODAE+ shared embedding. Each point represents a sample and is coloured according to tissue of origin; crosses denote TCGA patient tumour samples, and circles denote cancer cell lines. (F) Heatmap summarizing cancer-lineage classification performance across methods and ablated MODAE+ variants. Rows correspond to integration strategies or model variants, while columns report overall classification metrics followed by tissue-specific F1-scores for each cancer type. Overall accuracy and balanced accuracy quantify how well cancer lineage can be recovered from each embedding, whereas tissue-specific F1-scores indicate the extent to which individual tumour types remain distinguishable after integration. Higher values indicate stronger preservation of tumour-of-origin structure in the learned representation.

### Transferred sensitivity estimates predict patient survival in treated subsets

The clinical relevance of MODAE+ drug-sensitivity estimates was evaluated in treated TCGA patient subsets using treatment annotations and OS/PFI survival endpoints, as described in Materials and Methods. In total, 54 CTRP compounds could be matched to TCGA treatment records, of which 29 had at least one cancer type with more than five treated patients. After excluding non-evaluable analyses because of insufficient observations or lack of variation in predicted sensitivities, 27 drugs and 118 cancer–drug pairs remained evaluable for MODAE+ survival validation. The complete cancer–drug-specific survival validation results, including treated patient numbers, Harrell’s C-index, Kaplan–Meier log-rank p-values and Cox proportional hazards statistics for each method, cancer type, drug and survival endpoint, are provided in **Supplementary Table S6**.

For direct comparison with CODE-AE-ADV, analyses were restricted to cancer–drug– endpoint pairs evaluable for both methods, yielding 41 OS and 37 PFI comparisons on the same treated TCGA subsets. Table 1 summarizes the mean Harrell’s C-index and the proportions of comparisons with significant Kaplan–Meier log-rank and Cox proportional hazards associations for each method. MODAE+ showed stronger survival relevance for both endpoints. In this setting, MODAE+ showed stronger survival relevance for both OS and PFI. For OS, MODAE+ achieved a mean C-index of 0.684 compared with 0.506 for CODE-AE-ADV, with significant associations in 46.3% of pairs by Kaplan–Meier analysis and 41.5% by Cox analysis, compared with 2.4% and 4.9% for CODE-AE-ADV, respectively. For PFI, MODAE+ achieved a mean C-index of 0.617 compared with 0.549 for CODE-AE-ADV, with significant association rates of 32.4% by both Kaplan–Meier and Cox analyses, compared with 5.4% and 10.8% for CODE-AE-ADV. These results indicate that transferred MODAE+ sensitivity estimates retain treatment-associated survival information in TCGA patients.

**Table 1.** Paired survival validation of predicted drug sensitivities in treated TCGA patients. Survival associations were evaluated for overlapping cancer–drug–endpoint pairs that were evaluable for both MODAE+ and CODE-AE-ADV, enabling a direct paired comparison between methods on the same treated TCGA subsets. For each algorithm and survival endpoint, the table reports the mean Harrell’s C-index, the proportion of cancer–drug–endpoint pairs with a significant Kaplan–Meier log-rank test, the proportion with a significant Cox proportional hazards association, and the number of evaluable pairs tested. OS, overall survival; PFI, progression-free interval; KM, Kaplan–Meier; PH, proportional hazards.

| Algorithm | Survival target | Mean C-index | KM log-rank significant rate | Cox PH significant rate | Number of pairs tested |
|---|---|---|---|---|---|
| MODAE+ | OS | 0.684 | 0.463 | 0.415 | 41 |
| MODAE+ | PFI | 0.617 | 0.324 | 0.324 | 37 |
| CODE-AE-ADV | OS | 0.506 | 0.024 | 0.049 | 41 |
| CODE-AE-ADV | PFI | 0.549 | 0.054 | 0.108 | 37 |

Because treated TCGA subsets included drugs with distinct mechanisms of action, survival associations were further summarized by pharmacological class (**Figure 2**). Class-stratified survival analysis showed that the clinical relevance of MODAE+ sensitivity estimates was not uniformly distributed across pharmacological classes. In the paired benchmark, MODAE+ showed stronger class-level survival evidence than CODE-AE-ADV across several treatment classes, despite both methods being evaluated on the same cancer–drug– endpoint pairs. This difference was particularly evident for endocrine therapy, antimetabolites, topoisomerase/DNA-damage agents, microtubule/mitotic-spindle agents, and multi-RTK/kinase-targeted agents, whereas CODE-AE-ADV showed limited Cox-significant evidence across most classes. The all-evaluable MODAE+ analysis confirmed that the signal was not simply explained by the number of evaluated pairs: highly represented classes such as platinum agents showed limited Cox-significant evidence, while less represented classes such as endocrine therapy displayed stronger associations. These results suggest that MODAE+ captures treatment-relevant variation in a drug-class-dependent manner, with stronger survival evidence in classes where transcriptomic state is more directly linked to pharmacological response.

**Figure 2.**
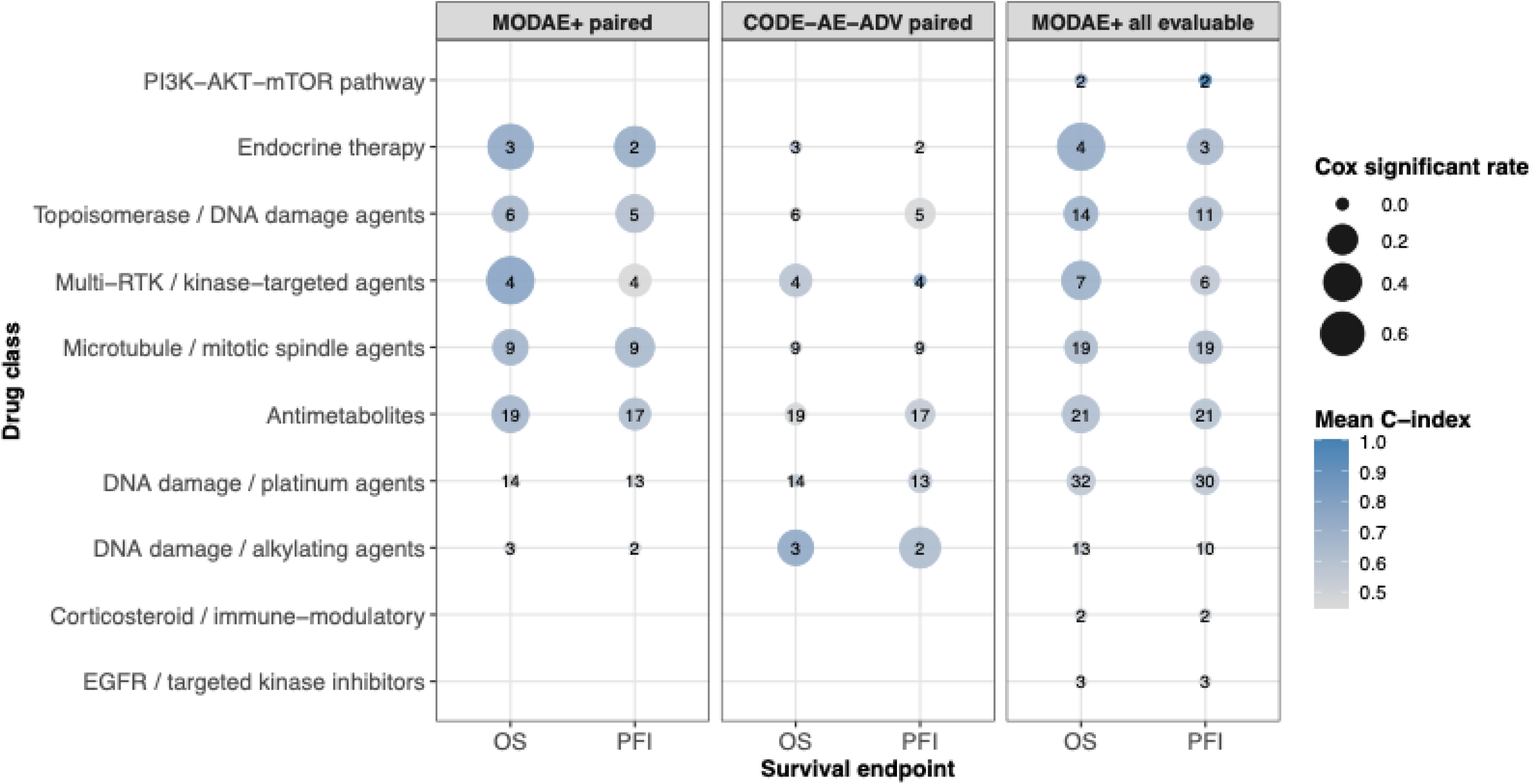
Drug-class-level survival evidence for transferred sensitivity predictions. Cancer–drug–endpoint survival associations were summarized by pharmacological class. The first two panels show the paired benchmark restricted to cancer–drug–endpoint pairs evaluable for both MODAE+ and CODE-AE-ADV, whereas the third panel shows all MODAE+ evaluable pairs. Bubble size indicates the proportion of pairs with significant Cox proportional hazards association, colour indicates the mean Harrell’s C-index, and the number inside each bubble indicates the number of evaluated cancer–drug–endpoint pairs. OS, overall survival; PFI, progression-free interval.

### Drug-target expression correlates with sensitivity estimate in patients

MODAE+ drug-sensitivity predictions for 9,808 TCGA patients were evaluated for biological relevance by comparing annotated drug-associated gene expression between predicted-sensitive and predicted-resistant groups. For this validation analysis, four annotation resources were examined, restricting analyses to drug–cancer combinations supported by Open Targets indication annotations (see Materials and Methods). **Table 2** summarizes the drug-associated expression differences meeting nominal p < 0.05 and |log□FC| > log□(1.5) across four annotation resources. Focusing on the lowest-p-value annotated gene for each drug, we identified significant differences for 34.9–64.4% of drugs when tumours were analysed together and for 29.6–48.6% of drug–cancer pairs when cancer types were analysed separately. Supporting associations were therefore observed both across tumours and within individual cancer types. PharmGKB yielded the highest rates in both settings, 64.4% and 48.6%, respectively, which may reflect its emphasis on pharmacogenomic determinants of drug response and systematic evidence curation (Gong *et al.*, 2021; Whirl Carrillo *et al.*, 2021). The complementary coverage of the four resources allowed us to examine intended molecular targets alongside genes involved in drug disposition and resistance. Individual drug–gene–cancer results are provided in **Supplementary Table S7**.

**Table 2.**
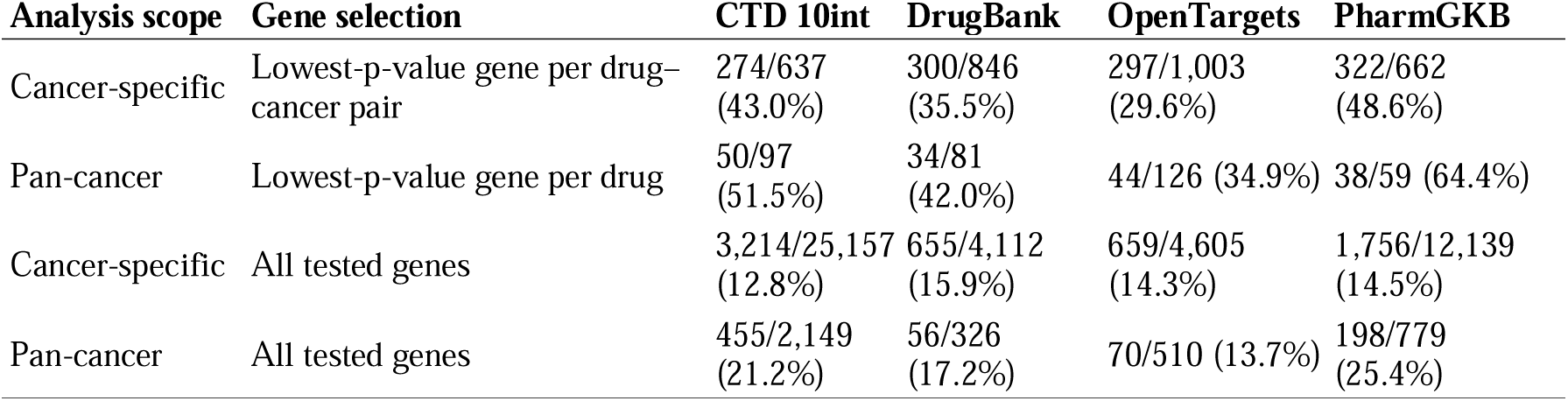
Drug-associated gene-expression differences between MODAE+ predicted-sensitive and predicted-resistant TCGA groups. Values indicate associations meeting both nominal p < 0.05 and |log FC| > log (1.5), divided by the number evaluated, with percentages in parentheses. Analyses used the indication-filtered subset. Minimum-p-value strategies selected one gene within each annotation source before applying both criteria.

| **Analysis scope** | **Gene selection** | **CTD 10int** | **DrugBank** | **OpenTargets** | **PharmGKB** |
|---|---|---|---|---|---|
| Cancer-specific | Lowest-p-value gene per drug–cancer pair | 274/637 (43.0%) | 300/846 (35.5%) | 297/1,003 (29.6%) | 322/662 (48.6%) |
| Pan-cancer | Lowest-p-value gene per drug | 50/97 (51.5%) | 34/81 (42.0%) | 44/126 (34.9%) | 38/59 (64.4%) |
| Cancer-specific | All tested genes | 3,214/25,157 (12.8%) | 655/4,112 (15.9%) | 659/4,605 (14.3%) | 1,756/12,139 (14.5%) |
| Pan-cancer | All tested genes | 455/2,149 (21.2%) | 56/326 (17.2%) | 70/510 (13.7%) | 198/779 (25.4%) |

Moreover, to assess whether transferred MODAE+ sensitivity predictions reflected biologically meaningful drug mechanisms, expression of curated drug target-associated genes was compared between predicted sensitive and resistant TCGA tumours across selected pharmacological classes (Figure 3).

**Figure 3.**
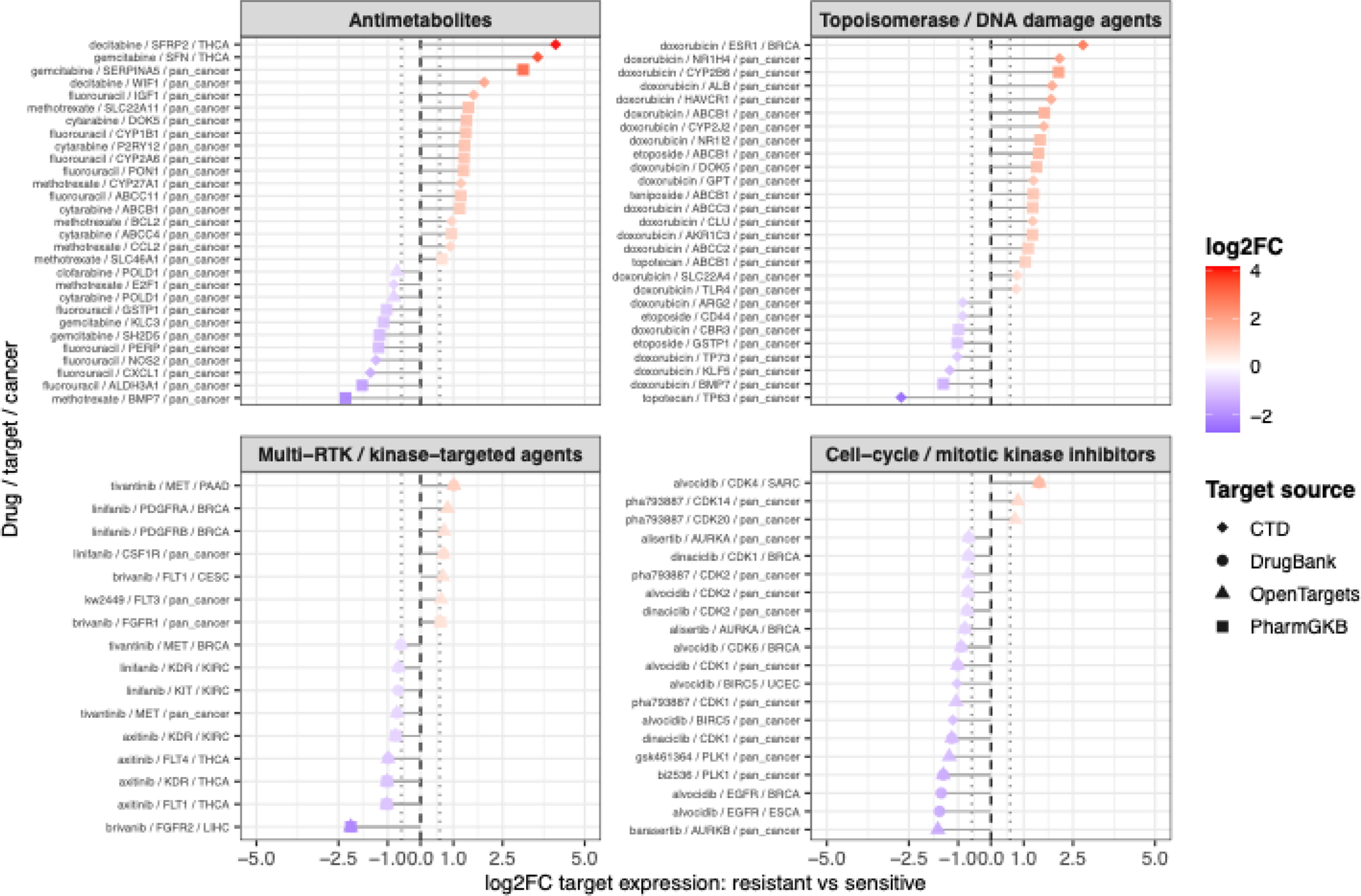
Drug-target expression differences between MODAE+ predicted sensitive and resistant TCGA patients. Top-ranked drug–target–cancer associations are shown for selected pharmacological classes. For each drug, TCGA patients were stratified into predicted sensitive and resistant groups according to MODAE+ sensitivity estimates, and expression of annotated drug target-associated genes was compared between groups using two-sided t-tests. Associations were restricted to the indication-filtered subset and required nominal p<0.05 and |*log_2_FC|*>*log*_2_(1.5). The top 30 associations per drug class were selected according to a combined statistical-evidence and effect-size score. Positive *log_2_FC* values indicate higher target-gene expression in predicted resistant patients, whereas negative values indicate higher expression in predicted sensitive patients. Point colour indicates *log_2_FC*, and point shape indicates the target-annotation source. The figure highlights drug-target expression patterns consistent with known mechanisms across antimetabolites, topoisomerase/DNA damage agents, multi-RTK/kinase-targeted agents, and cell-cycle/mitotic kinase inhibitors.

Stratifying these associations by drug class revealed a clear distinction between cytotoxic chemotherapies and mechanism-targeted agents. For cytotoxic chemotherapies, including antimetabolites and topoisomerase/DNA-damage agents, the strongest associations were frequently observed for genes involved in drug transport, metabolism, detoxification, and resistance rather than for the intended molecular targets. ATP-binding cassette transporters such as *ABCB1*, *ABCC1–4*, and *ABCG2* appeared across multiple chemotherapy agents, together with drug-metabolizing or detoxification genes including *CYP1B1*, *CYP2A6*, *CYP2B6*, *GSTP1*, *AKR1C3*, *CBR3*, and *ALDH3A1*. This pattern is consistent with the broad mechanisms of cytotoxic chemotherapy, where response and resistance are often mediated by altered drug accumulation, metabolism, or detoxification. By contrast, mechanism-targeted agents showed more direct target-related expression patterns. Multi-RTK and kinase-targeted inhibitors were associated with annotated receptor or kinase targets, including *KDR*, *FLT1*, and *FLT4* for axitinib; *PDGFRA*, *PDGFRB*, *KIT*, and *CSF1R* for linifanib; *FGFR1* and *FGFR2* for brivanib; and *FLT3* for KW-2449. Similarly, cell-cycle and mitotic-kinase inhibitors recovered expected target families, including *PLK1* for BI2536 and GSK461364, *AURKA/AURKB* for alisertib and barasertib, and multiple CDK genes for dinaciclib and alvocidib. The same pattern was supported by additional pharmacological classes shown in Supplementary Figure S3. MAPK pathway inhibitors recovered several pathway-associated genes, including *BRAF*, *KRAS*, *HRAS*, *NRAS*, and *MAP2K2*, together with apoptosis-related downstream markers such as *CASP3* and *CASP8*. Epigenetic and chromatin-regulating drugs showed associations with HDAC-related genes, including *HDAC1*, *HDAC2*, *HDAC7*, and *HDAC9*, together with additional context-specific transcriptional response genes. PARP/DNA repair agents showed several olaparib-associated signals involving *BRCA1*, *BRCA2*, and *PARP3*, whereas platinum agents and microtubule/mitotic-spindle drugs again highlighted transporter, detoxification, and resistance-related genes, including *GSTP1*, *ABCG2*, *ABCC2*, *ABCB1*, and taxane-associated genes such as *MAPT* and *TUBB3*.

### External response signatures support breast cancer chemotherapy concordance

As an additional validation, we examined whether MODAE+ predictions agreed with pretreatment expression signatures associated with clinical response in external breast cancer cohorts. To this end, signatures derived from 13 GEO accessions were applied to TCGA breast cancer patients who had received paclitaxel or doxorubicin, assessing whether higher predicted sensitivity corresponded to expression profiles resembling those of clinical responders. We first assessed the signatures’ ability to distinguish responders from nonresponders within their original clinical cohorts (**Supplementary Figure S4; Supplementary Table S10**). Top-200 signatures achieved cross-validated AUCs above 0.50 in 35 of 40 source groups (87.5%), including nine with AUC ≥ 0.70. Thus, most signatures showed response discrimination in the expected direction, although its strength varied across cohorts. Notably, reducing signature size did not generally improve performance. Median AUCs were 0.636 for top-200 signatures, compared with 0.624 for top-50 and 0.611 for top-10 signatures. Smaller signatures outperformed the corresponding top-200 signature in only 14 of 40 comparisons for top-50 and 17 of 40 for top-10. Pathway restriction nevertheless improved discrimination in several source groups: spindle, DNA-repair and E2F signatures exceeded the corresponding top-200 AUC in 16 of 36, 16 of 38 and 16 of 40 comparisons, respectively. For example, DNA-repair restriction increased AUC from 0.63 to 0.71 in GSE123845, while E2F restriction increased AUC from 0.64 to 0.73 in GSE20271 Test 1. These context-dependent gains suggest that biologically focused gene selection can capture response information more effectively than differential-expression ranking alone in some treatment settings. However, no pathway improved performance overall, indicating that pathway-based biomarkers require validation within the relevant treatment and tumour context. We next assessed concordance between external response-signature scores and MODAE+ predictions in TCGA breast cancer patients treated with paclitaxel or doxorubicin (Figure 4, left panel; Supplementary Table S11).

**Figure 4.**
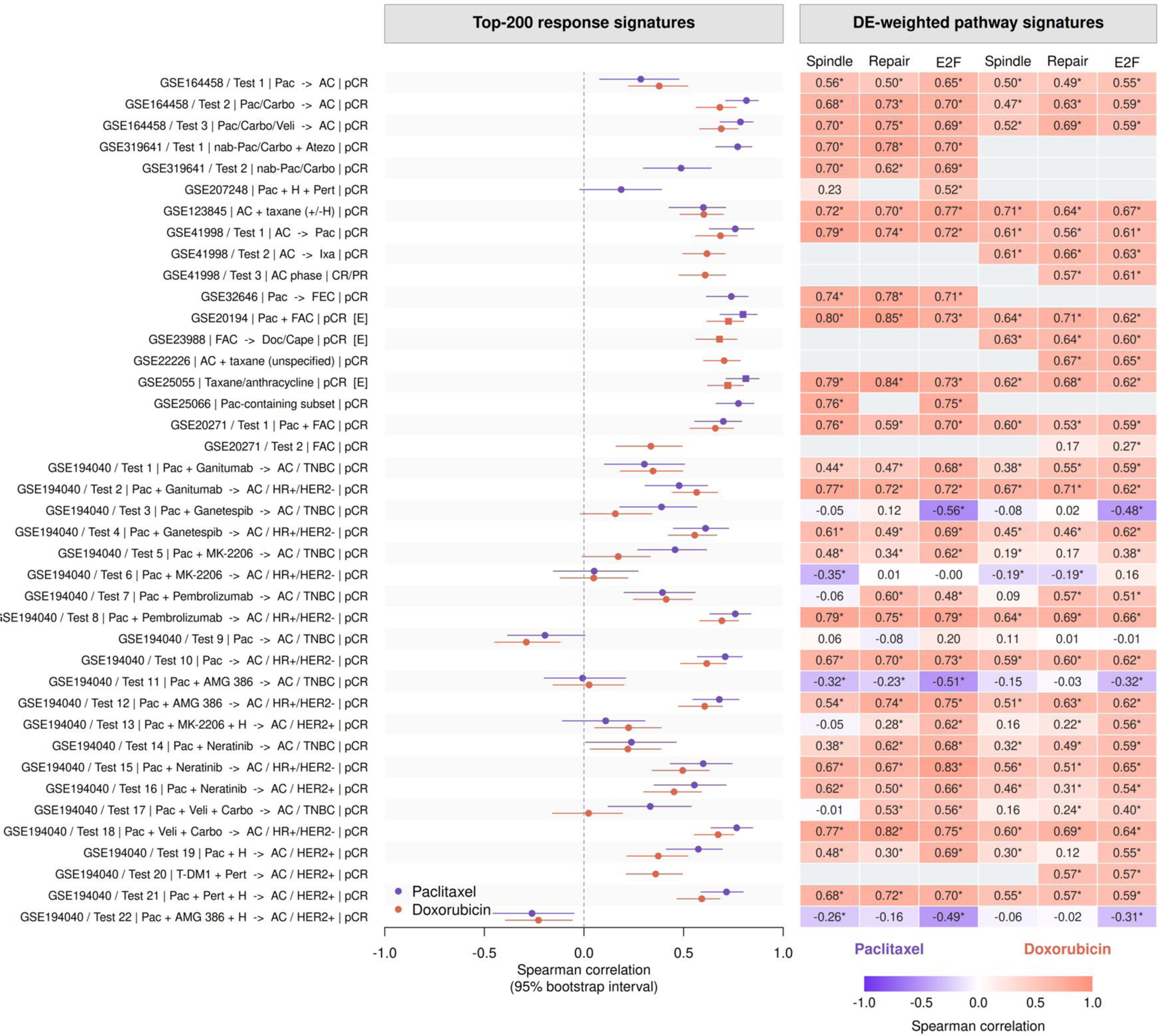
Concordance between MODAE+ predictions and clinical response-derived signatures in breast cancer. Pretreatment responder–nonresponder signatures from external cohorts were scored in TCGA breast tumours exposed to paclitaxel or doxorubicin. Left: Spearman correlations for top-200 signatures across 69 comparisons from 40 source groups and 13 GEO accessions; bars indicate 95% bootstrap confidence intervals. Purple denotes paclitaxel and coral doxorubicin. Right: Correlations for DE-weighted Hallmark spindle, DNA-repair and E2F signatures; purple indicates negative and coral positive correlations, with unavailable results in grey. All correlations use original MODAE+ predictions in held-out patients. Asterisks denote q < 0.05 across 1,205 comparisons. [E[E] labels and square markers identify exploratory comparisons in which treatment metadata did not permit confident attribution of the source signature to the individual drug evaluated; this designation is independent of statistical significance. Regimens and response definitions are detailed in Supplementary Table S8. pCR, pathological complete response; CR/PR, complete or partial clinical response.

Of the 69 top-200 signature–drug comparisons, 64 showed positive correlations, 56 were positive at FDR-adjusted q < 0.05, and 37 reached Spearman’s ρ ≥ 0.50. Thus, higher predicted sensitivity generally corresponded to more responder-like expression profiles, although the strength of agreement varied across source treatment arms and receptor subgroups. Subsequently, we examined whether restricting signatures to response-associated genes within selected pathways strengthened concordance with MODAE+ predictions (**Figure 4, right panel; Supplementary Table S11**). E2F-restricted signatures improved correlations in 40 of 69 matched comparisons (58%), with a median increase in Spearman’s ρ of 0.021. DNA-repair signatures improved 35 of 67 comparisons (52%), while mitotic-spindle signatures improved 28 of 65 (43%). Several pathway-restricted scores achieved correlations above 0.70, indicating that strong agreement could be recovered using genes belonging to defined biological programmes.

Finally, we summarized concordance by drug and source platform (Table 3). Positive, statistically significant correlations occurred in 28 of 34 paclitaxel comparisons and 28 of 35 doxorubicin comparisons, including 10 of 11 RNA-seq-derived and 46 of 58 microarray-derived comparisons. These results show that agreement with MODAE+ predictions extended across both drugs and expression platforms.

**Table 3.** Summary of concordance between external breast cancer response signatures and MODAE+ predictions. Counts of top-200 signature–drug comparisons with positive Spearman correlations, positive correlations at Benjamini– Hochberg q < 0.05, and correlations ≥ 0.50, grouped by drug and source expression platform. Drug and platform rows summarize the same 69 comparisons and are not additive.

| **Group** | **Total comparisons** | **Positive ρ** | **Positive with q < 0.05** | **ρ ≥ 0.50** |
|---|---|---|---|---|
| All | 69 | 64 | 56 | 37 |
| Paclitaxel | 34 | 31 | 28 | 19 |
| Doxorubicin | 35 | 33 | 28 | 18 |
| Microarray-derived | 58 | 53 | 46 | 30 |
| RNA-seq-derived | 11 | 11 | 10 | 7 |

Combining eligible external signatures into patient-level consensus scores yielded Spearman correlations of 0.791 for paclitaxel (95% CI, 0.685–0.861; 83 evaluation patients) and 0.724 for doxorubicin (0.620–0.798; 130 evaluation patients), incorporating nine and six GEO accessions, respectively. The equal-weight mean correlation across the two drugs was 0.758 (95% CI, 0.664–0.817), supporting strong concordance between MODAE+ predictions and response-associated expression patterns aggregated across external studies.

## Conclusions

MODAE+ addresses a central limitation in pharmacogenomic transfer learning: aligning cancer cell lines and patient tumours while preserving biologically meaningful cancer-lineage structure. By combining adversarial domain alignment, deconfounding, survival supervision, drug-response prediction, cancer-lineage classification and adaptive multi-objective optimization, MODAE+ learned a shared latent representation that preserved tumour-of-origin organization while reducing cell-line–tumour domain separation. Ablation analyses indicated that this balance depended on the full multi-objective architecture.

The transferred drug-sensitivity estimates were supported by three complementary validation approaches. Among TCGA patients with documented receipt of the evaluated drugs, MODAE+ predictions showed stronger associations with survival than CODE-AE-ADV, supporting their relevance to treatment-associated patient variation. Analyses using curated drug–gene annotations also recovered expected expression patterns: differences between predicted-sensitive and predicted-resistant tumours involved intended molecular targets for targeted agents and predominantly transport, metabolism, detoxification and resistance-related programmes for classical chemotherapies.

Additional validation using clinical-response signatures from 13 external breast cancer GEO accessions showed predominantly positive concordance with paclitaxel and doxorubicin predictions across RNA-seq- and microarray-derived signatures. Patient-level consensus scores achieved correlations of 0.791 and 0.724, respectively, with a mean of 0.758 across the two drugs. The consistency across external signatures and strong consensus agreement strengthen confidence that MODAE+ captures expression patterns associated with chemotherapy response. This post hoc analysis remains specific to the selected breast cancer treatments, with overlapping cohorts and combination regimens limiting the independence and drug specificity of the evidence. Direct clinical predictive accuracy therefore requires further evaluation against observed treatment outcomes.

Overall, the complementary validation results support the robustness of MODAE+ predictions in the evaluated settings, although their reliability may vary across cancer types and drugs according to the representation and quality of the corresponding cell-line expression and drug-response data in CCLE and CTRP.

## Key Points

- MODAE+ is a pan-cancer multi-objective domain-adaptation framework that transfers pharmacogenomic drug-response information from cancer cell lines to patient tumours by jointly optimizing domain alignment, deconfounding, drug-response prediction, survival modelling and cancer-lineage classification, with objectives adaptively balanced by FAMO.
- MODAE+ improved cell-line–tumour alignment while preserving tumour-of-origin organization, recovering cancer-lineage structure more faithfully than CODE-AE-ADV in the evaluated benchmark.
- In treated TCGA subsets, MODAE+ sensitivity estimates were associated with survival more frequently than CODE-AE-ADV across matching cancer–drug–endpoint comparisons, including an overall-survival Cox-significant rate of **41.5% versus 4.9%**.
- Drug-associated expression analyses provided biological support for the predictions. Top-ranked associations included intended molecular targets for targeted therapies and genes involved in transport, metabolism, detoxification and resistance for classical chemotherapies.
- External clinical-response signatures from **13 breast cancer GEO accessions** showed predominantly positive concordance with MODAE+ predictions across **69 comparisons**, with consensus correlations of **0.791 for paclitaxel** and **0.724 for doxorubicin**, providing complementary expression-based validation of predicted drug sensitivities.

## Acknowledgements

The computational analyses were performed on servers provided by the Bioinformatics Center of the University of Eastern Finland.

## Author Contributions

T.J.R.: Writing – review & editing, Writing, Visualization, Software, Methodology, Investigation, Formal analysis, Data curation, V.F.: Writing (original draft), review & editing, Supervision, Project Administration, Investigation, Formal analysis, and Funding acquisition.

## DATA AVAILABILITY

The datasets generated and/or analyzed during the present study are publicly available in the Zenodo repository at https://doi.org/10.5281/zenodo.21808125.

## CODE AVAILABILITY

All source code and scripts required to reproduce the results and analyses presented in this study are available on GitHub at https://github.org/UEFBiomedicalInformaticsLab/MODAEplus.

## SUPPLEMENTARY FIGURE LEGENDS

**Supplementary Figure S1.**
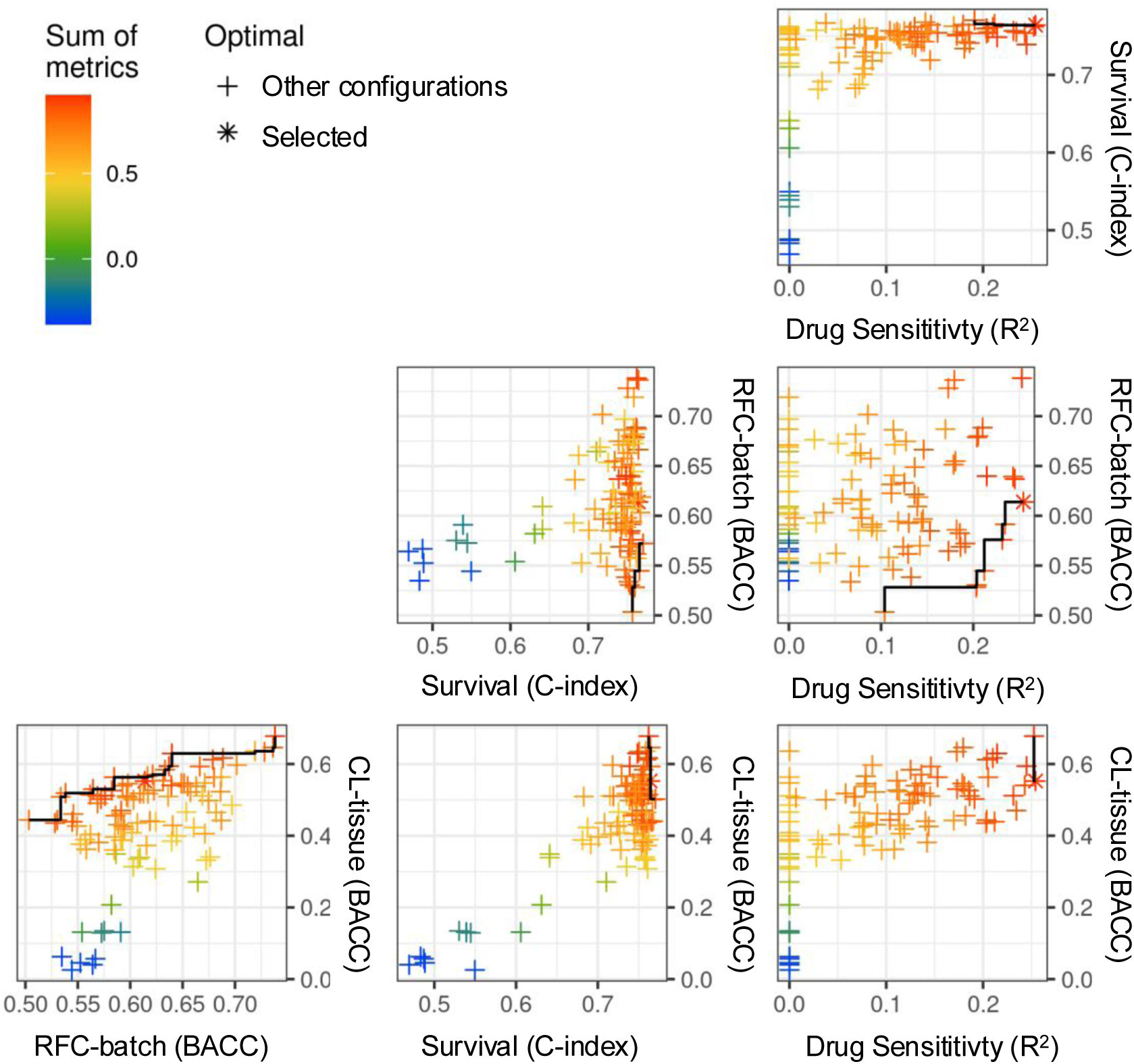
Pairwise relationships between optimization objectives across random-search configurations. Each point represents one randomly sampled MODAE+ hyperparameter configuration evaluated during cross-validation. Axes show pairwise comparisons between the objectives used for model selection, including patient survival C-index, drug sensitivity, domain prediction balanced accuracy based on the batch random forest classifier, and cancer lineage classification balanced accuracy for cell-line samples. Point colour indicates the composite optimization score, defined as the weighted sum of the selected performance metrics. The final selected configuration, used for all downstream analyses, is marked with an asterisk, while all other evaluated configurations are shown with crosses. The selected model achieved a favourable trade-off across objectives, combining strong survival performance, measurable drug-response prediction, preserved cancer-lineage classification, and improved cross-domain alignment.

**Supplementary Figure S2.**
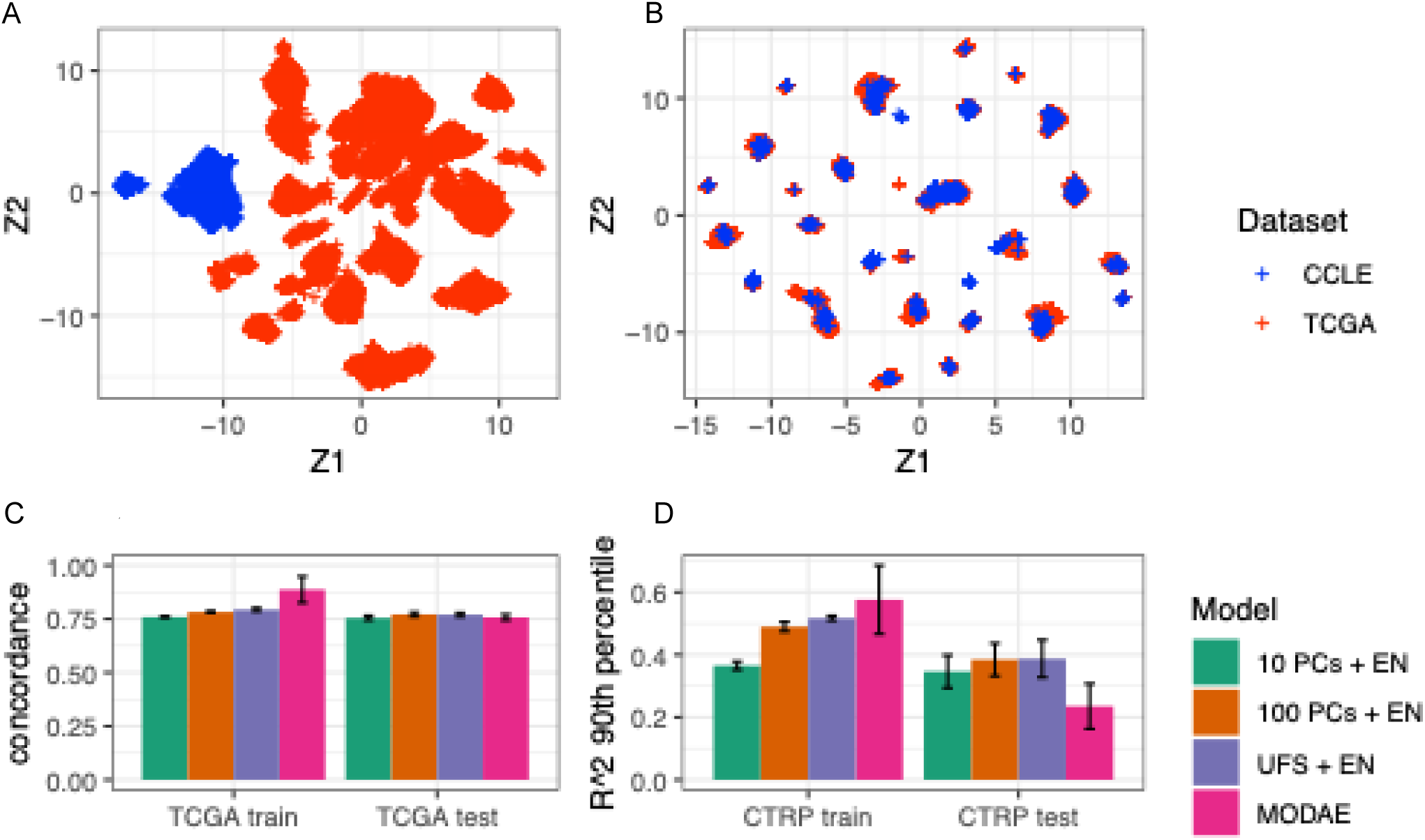
Test-set performance of the selected MODAE+ model. Performance of the final MODAE+ configuration selected from the random-search optimization. **(A)** Latent-space projection before domain adaptation showing clear separation between CCLE cell lines and TCGA tumour samples. **(B)** Latent-space projection produced by the selected MODAE+ model, showing improved mixing of CCLE and TCGA samples while retaining structured cancer-specific organization. **(C)** Survival prediction performance in TCGA training and test sets, reported as concordance index and compared with elastic-net baseline models. **(D)** Drug-response prediction performance in CTRP training and test sets, reported as for the 90th percentile of drug-sensitivity predictions and compared with elastic-net baseline models. Error bars indicate variability across cross-validation folds. Overall, the selected MODAE+ model improved cross-domain alignment and maintained competitive survival prediction performance, while held-out CTRP drug-response prediction showed weaker generalization, consistent with the known difficulty of transferring drug-response models to independent test data.

**Supplementary Figure S3.**
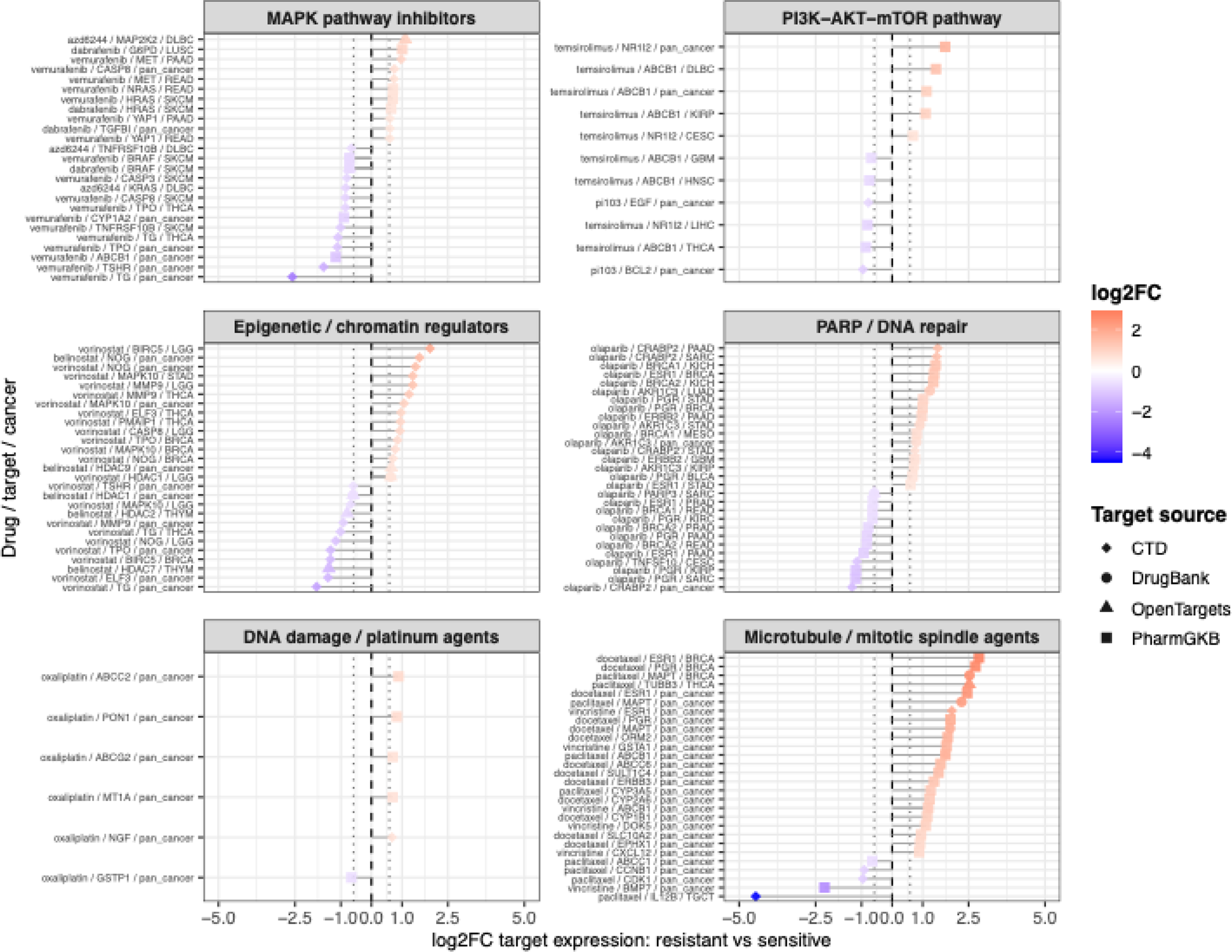
Additional drug-target expression differences between MODAE+ predicted sensitive and resistant TCGA patients. Top-ranked drug–target– cancer associations are shown for additional pharmacological classes not included in the main figure. Patients were stratified into predicted sensitive and resistant groups according to MODAE+ drug-sensitivity estimates, and expression differences of annotated drug target-associated genes were evaluated using two-sided t-tests. Associations were restricted to the indication-filtered subset and required nominal p <0.05 and |*log_2_FC|*>*log*_2_(1.5). Up to 30 top-ranked associations per drug class are shown, selected using a combined score based on statistical evidence and effect size. Positive *log_2_FC*values indicate higher target-gene expression in predicted resistant patients, whereas negative values indicate higher expression in predicted sensitive patients. Point colour indicates, and point shape indicates the target-annotation source. The figure provides additional examples of drug-target expression patterns across MAPK pathway inhibitors, PI3K–AKT–mTOR pathway inhibitors, epigenetic/chromatin regulators, PARP/DNA repair agents, DNA damage/platinum agents, and microtubule/mitotic spindle agents.

**Supplementary Figure S4.**
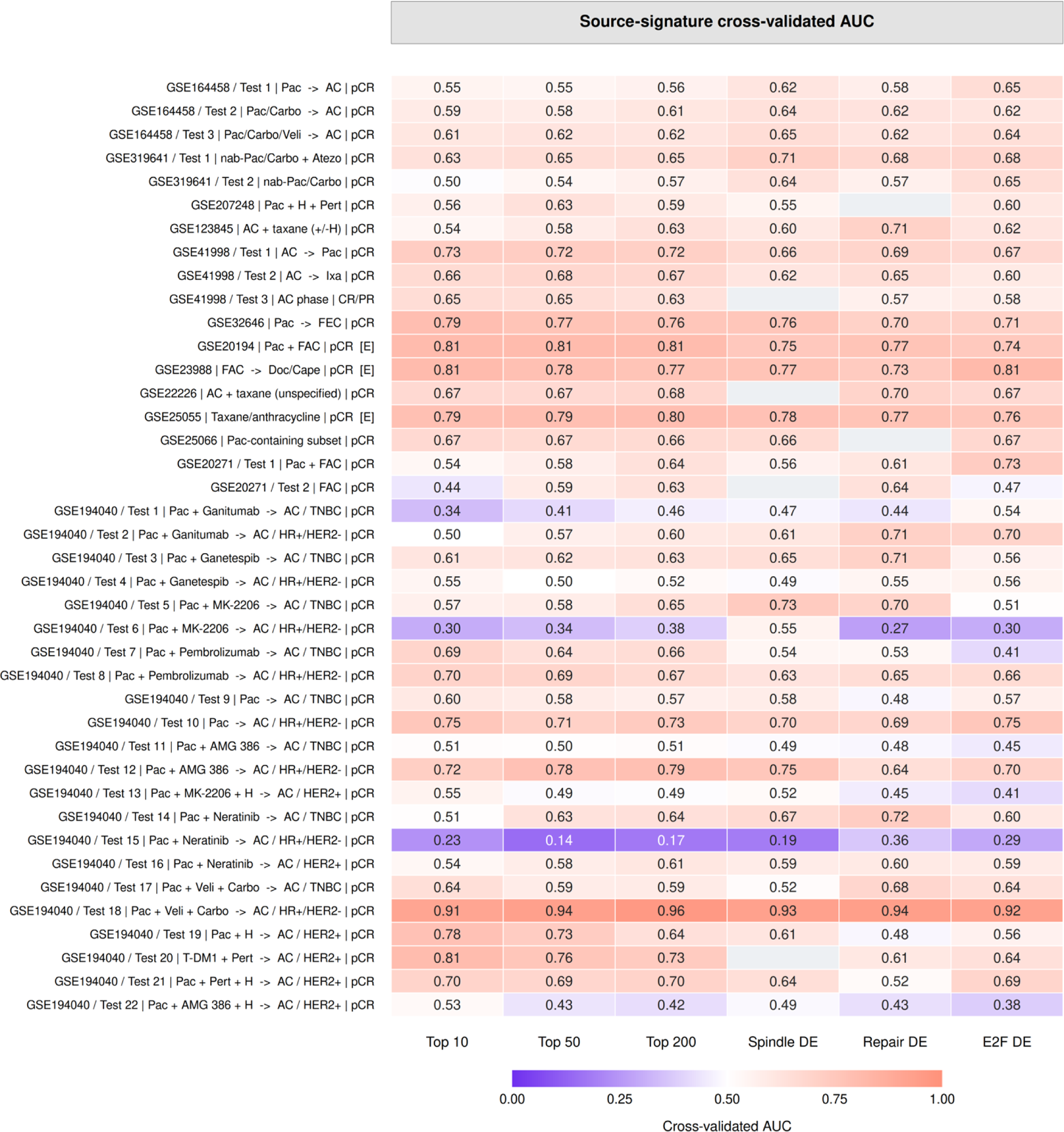
Ability of the external gene-expression signatures to distinguish treatment responders from nonresponders. Signatures were constructed from pretreatment expression data using the top 10, 50 or 200 differentially expressed genes, or genes restricted to the indicated pathways. Performance was evaluated by repeated patient-level cross-validation, with gene selection, weighting and standardization performed within each training fold before scoring held-out patients. Cells show mean receiver operating characteristic AUC: 0.5 indicates chance discrimination and higher values indicate better discrimination. Below-chance values are retained; grey cells indicate unavailable results. Rows correspond to Figure 4. These results evaluate the response-derived signatures, not MODAE+ predictions directly. Further details are provided in Materials and Methods.

## SUPPLEMENTARY TABLE LEGENDS

**Supplementary Table S1. TCGA cancer lineage annotations used for MODAE+ lineage supervision.** The table reports the mapping of TCGA tumour samples to harmonized OncoTree Level 1 cancer lineage categories, together with the corresponding TCGA tumour type labels.

**Supplementary Table S2. CCLE cancer cell-line lineage annotations used for MODAE+ lineage supervision.** The table reports the mapping of CCLE/DepMap cell lines to harmonized OncoTree Level 1 cancer lineage categories, including the corresponding primary lineage annotation when available.

**Supplementary Table S3. Hyperparameter search space used for MODAE+ tuning.** The table reports the model architecture, optimization, regularization, and FAMO-related hyperparameters explored during random search, including the corresponding candidate values or value ranges.

**Supplementary Table S4. Cross-validation performance summary for MODAE+ hyperparameter configurations.** The table reports training and test performance metrics for each randomly sampled hyperparameter configuration evaluated during model selection, including reconstruction, survival prediction, drug-response prediction, domain-alignment, and cancer-lineage classification metrics where applicable.

**Supplementary Table S5. Drug-to-pharmacological-class mapping used for class-stratified analyses.** The table reports the pharmacological class assigned to each drug included in the drug-target expression and/or treatment-specific survival analyses. For each drug, the table indicates the assigned mechanism-based class and whether the drug was used in the drug-target expression analysis, the survival validation analysis, or both. This mapping was used to stratify drug-target expression associations and survival-validation results by pharmacological class.

**Supplementary Table S6**. **Cancer–drug-specific survival validation of predicted drug sensitivities in treated TCGA patient subsets.** Each row corresponds to one method, cancer type, drug, and survival endpoint combination. The table reports the number of treated patients evaluated, Harrell’s concordance index, Kaplan–Meier log-rank p-value comparing predicted sensitive and resistant groups, and Cox proportional hazards statistics evaluating predicted sensitivity as a continuous covariate. Overall survival and progression-free interval endpoints are reported when evaluable.

**Supplementary Table S7**. **Drug-target expression associations between MODAE+ predicted sensitive and resistant TCGA tumours.** The table reports the complete set of drug–target gene expression comparisons used to assess whether MODAE+ predicted drug sensitivity groups were associated with biologically meaningful target-related molecular differences. For each cancer type, drug, and curated target-associated gene, TCGA tumours were stratified into predicted sensitive and resistant groups according to the MODAE+ sensitivity estimate for that drug. The table reports the mean gene expression in the predicted sensitive and resistant groups, the number of samples in each group, the nominal two-sided t-test p-value, the log fold change between resistant and sensitive groups, and the target-annotation source. Positive log FC values indicate higher target-gene expression in predicted resistant tumours, whereas negative values indicate higher expression in predicted sensitive tumours.

**Supplementary Table S8. External breast cancer source groups.** GEO accession, expression platform, treatment regimen, receptor subtype, response definitions, responder and nonresponder sample sizes, and Figure 4 labels for 40 source groups from 13 accessions.

**Supplementary Table S9. Expression data sources and preprocessing.** Processed matrix download URLs, MD5 checksums, expression units, transformations and gene-mapping information for the 13 GEO accessions.

**Supplementary Table S10. Clinical-response discrimination of external signatures.** Mean ROC AUCs from five repeats of stratified cross-validation, with up to five folds per repeat, and response-group sample sizes for top-10, top-50, top-200 and pathway-restricted signatures. These AUCs evaluate source-cohort signatures, not MODAE+ predictive accuracy.

**Supplementary Table S11. Concordance between external signatures and MODAE+ predictions in TCGA breast cancer.** Spearman correlations, bootstrap confidence intervals and Benjamini–Hochberg q values for evaluable paclitaxel and doxorubicin comparisons. Results include original and PC1-subtracted predictions and the selected result; original predictions were retained for both drugs in Figure 4. Q values were calculated across the broader set of 1,205 comparisons.

## SUPPLEMENTARY FILES

**Supplementary File S1. Mathematical formulation and training algorithm of MODAE+.** Detailed model equations, loss functions and joint-training pseudocode, including adversarial alignment, deconfounding, drug-response and survival prediction, cancer-lineage classification and FAMO-based multi-objective optimization.

