## Supplementary Files Figures And Tables for "Multi-objective domain adaptation for pan-cancer transfer of pharmacogenomic drug response to patient tumours": Supplementary Figure S2.pdf

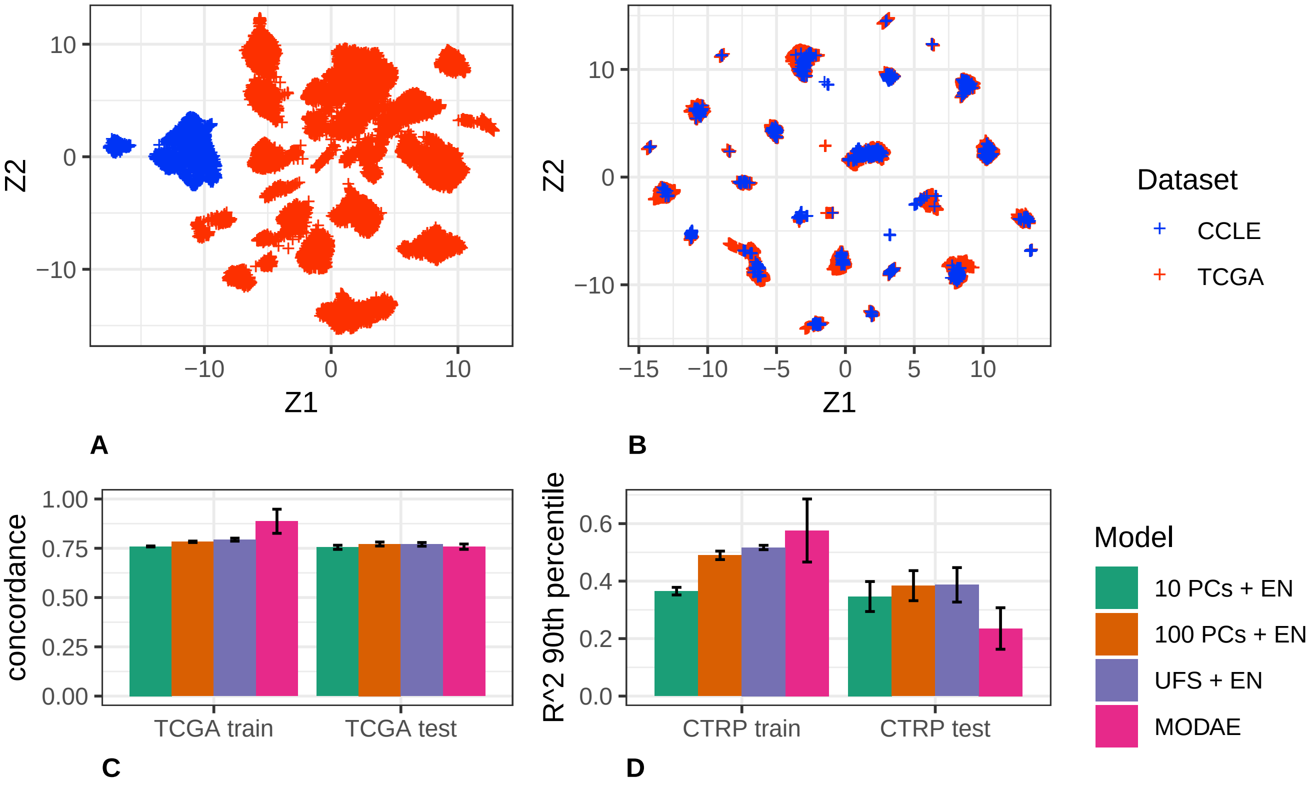

10
0
−10
Z2
−10
0
10
Z1
A
10
0
−10
Z2
−15
−10
−5
0
5
10
Z1
B
Dataset
CCLE
TCGA
1.00
0.75
0.50
0.25
0.00
concordance
TCGA train
TCGA test
C
0.6
0.4
0.2
0.0
R^2 90th percentile
CTRP train
CTRP test
D
Model
10 PCs + EN
100 PCs + EN
UFS + EN
MODAE
