## Supplementary Files Figures And Tables for "Multi-objective domain adaptation for pan-cancer transfer of pharmacogenomic drug response to patient tumours": Supplementary Figure S3.pdf

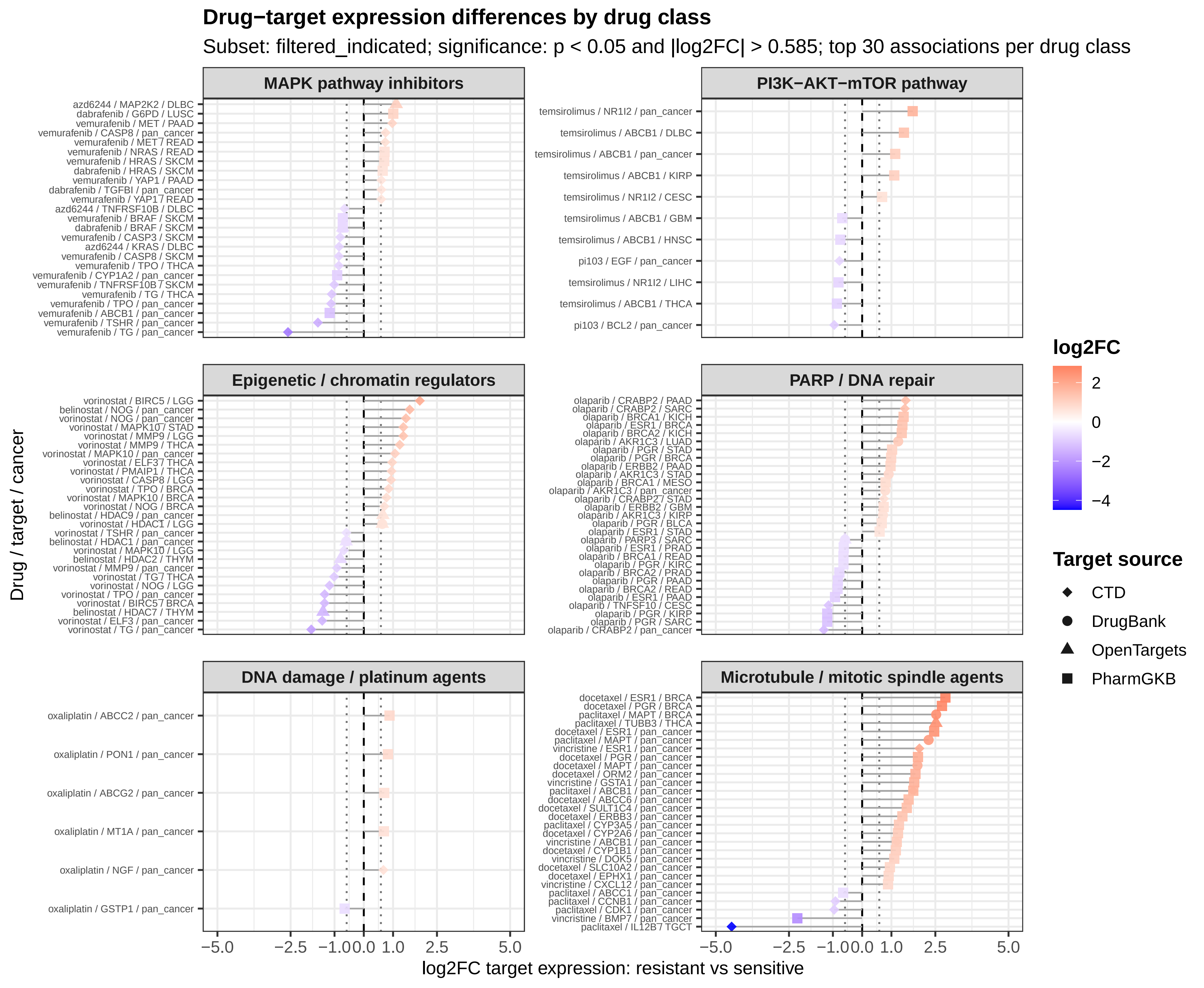

Drug−target expression differences by drug class
Subset: filtered_indicated; significance: p < 0.05 and |log2FC| > 0.585; top 30 associations per drug class
MAPK pathway inhibitors
azd6244 / MAP2K2 / DLBC
dabrafenib / G6PD / LUSC
vemurafenib / MET / PAAD
vemurafenib / CASP8 / pan_cancer
vemurafenib / MET / READ
vemurafenib / NRAS / READ
vemurafenib / HRAS / SKCM
dabrafenib / HRAS / SKCM
vemurafenib / YAP1 / PAAD
dabrafenib / TGFBI / pan_cancer
vemurafenib / YAP1 / READ
azd6244 / TNFRSF10B / DLBC
vemurafenib / BRAF / SKCM
dabrafenib / BRAF / SKCM
vemurafenib / CASP3 / SKCM
azd6244 / KRAS / DLBC
vemurafenib / CASP8 / SKCM
vemurafenib / TPO / THCA
vemurafenib / CYP1A2 / pan_cancer
vemurafenib / TNFRSF10B / SKCM
vemurafenib / TG / THCA
vemurafenib / TPO / pan_cancer
vemurafenib / ABCB1 / pan_cancer
vemurafenib / TSHR / pan_cancer
vemurafenib / TG / pan_cancer
PI3K−AKT−mTOR pathway
temsirolimus / NR1I2 / pan_cancer
temsirolimus / ABCB1 / DLBC
temsirolimus / ABCB1 / pan_cancer
temsirolimus / ABCB1 / KIRP
temsirolimus / NR1I2 / CESC
temsirolimus / ABCB1 / GBM
temsirolimus / ABCB1 / HNSC
pi103 / EGF / pan_cancer
temsirolimus / NR1I2 / LIHC
temsirolimus / ABCB1 / THCA
pi103 / BCL2 / pan_cancer
Epigenetic / chromatin regulators
vorinostat / BIRC5 / LGG
belinostat / NOG / pan_cancer
vorinostat / NOG / pan_cancer
vorinostat / MAPK10 / STAD
vorinostat / MMP9 / LGG
vorinostat / MMP9 / THCA
vorinostat / MAPK10 / pan_cancer
vorinostat / ELF3 / THCA
vorinostat / PMAIP1 / THCA
vorinostat / CASP8 / LGG
vorinostat / TPO / BRCA
vorinostat / MAPK10 / BRCA
vorinostat / NOG / BRCA
belinostat / HDAC9 / pan_cancer
vorinostat / HDAC1 / LGG
vorinostat / TSHR / pan_cancer
belinostat / HDAC1 / pan_cancer
vorinostat / MAPK10 / LGG
belinostat / HDAC2 / THYM
vorinostat / MMP9 / pan_cancer
vorinostat / TG / THCA
vorinostat / NOG / LGG
vorinostat / TPO / pan_cancer
vorinostat / BIRC5 / BRCA
belinostat / HDAC7 / THYM
vorinostat / ELF3 / pan_cancer
vorinostat / TG / pan_cancer
PARP / DNA repair
olaparib / CRABP2 / PAAD
olaparib / CRABP2 / SARC
olaparib / BRCA1 / KICH
olaparib / ESR1 / BRCA
olaparib / BRCA2 / KICH
olaparib / AKR1C3 / LUAD
olaparib / PGR / STAD
olaparib / PGR / BRCA
olaparib / ERBB2 / PAAD
olaparib / AKR1C3 / STAD
olaparib / BRCA1 / MESO
olaparib / AKR1C3 / pan_cancer
olaparib / CRABP2 / STAD
olaparib / ERBB2 / GBM
olaparib / AKR1C3 / KIRP
olaparib / PGR / BLCA
olaparib / ESR1 / STAD
olaparib / PARP3 / SARC
olaparib / ESR1 / PRAD
olaparib / BRCA1 / READ
olaparib / PGR / KIRC
olaparib / BRCA2 / PRAD
olaparib / PGR / PAAD
olaparib / BRCA2 / READ
olaparib / ESR1 / PAAD
olaparib / TNFSF10 / CESC
olaparib / PGR / KIRP
olaparib / PGR / SARC
olaparib / CRABP2 / pan_cancer
DNA damage / platinum agents
oxaliplatin / ABCC2 / pan_cancer
oxaliplatin / PON1 / pan_cancer
oxaliplatin / ABCG2 / pan_cancer
oxaliplatin / MT1A / pan_cancer
oxaliplatin / NGF / pan_cancer
oxaliplatin / GSTP1 / pan_cancer
Microtubule / mitotic spindle agents
docetaxel / ESR1 / BRCA
docetaxel / PGR / BRCA
paclitaxel / MAPT / BRCA
paclitaxel / TUBB3 / THCA
docetaxel / ESR1 / pan_cancer
paclitaxel / MAPT / pan_cancer
vincristine / ESR1 / pan_cancer
docetaxel / PGR / pan_cancer
docetaxel / MAPT / pan_cancer
docetaxel / ORM2 / pan_cancer
vincristine / GSTA1 / pan_cancer
paclitaxel / ABCB1 / pan_cancer
docetaxel / ABCC6 / pan_cancer
docetaxel / SULT1C4 / pan_cancer
docetaxel / ERBB3 / pan_cancer
paclitaxel / CYP3A5 / pan_cancer
docetaxel / CYP2A6 / pan_cancer
vincristine / ABCB1 / pan_cancer
docetaxel / CYP1B1 / pan_cancer
vincristine / DOK5 / pan_cancer
docetaxel / SLC10A2 / pan_cancer
docetaxel / EPHX1 / pan_cancer
vincristine / CXCL12 / pan_cancer
paclitaxel / ABCC1 / pan_cancer
paclitaxel / CCNB1 / pan_cancer
paclitaxel / CDK1 / pan_cancer
vincristine / BMP7 / pan_cancer
paclitaxel / IL12B / TGCT
Drug / target / cancer
−5.0 −2.5 −1.0 0.0 1.0 2.5 5.0
log2FC target expression: resistant vs sensitive
log2FC
2
0
−2
−4
Target source
CTD
DrugBank
OpenTargets
PharmGKB
