## Supplementary Files Figures And Tables for "Multi-objective domain adaptation for pan-cancer transfer of pharmacogenomic drug response to patient tumours": Supplementary Figure S4.pdf

### Supplementary Figure S4 | Source−signature cross−validated AUC

| | Top 10 | Top 50 | Top 200 | Spindle DE | Repair DE | E2F DE |
|---|---|---|---|---|---|---|
| GSE164458 / Test 1 \| Pac −> AC \| pCR | 0.55 | 0.55 | 0.56 | 0.62 | 0.58 | 0.65 |
| GSE164458 / Test 2 \| Pac/Carbo −> AC \| pCR | 0.59 | 0.58 | 0.61 | 0.64 | 0.62 | 0.62 |
| GSE164458 / Test 3 \| Pac/Carbo/Veli −> AC \| pCR | 0.61 | 0.62 | 0.62 | 0.65 | 0.62 | 0.64 |
| GSE319641 / Test 1 \| nab−Pac/Carbo + Atezo \| pCR | 0.63 | 0.65 | 0.65 | 0.71 | 0.68 | 0.68 |
| GSE319641 / Test 2 \| nab−Pac/Carbo \| pCR | 0.50 | 0.54 | 0.57 | 0.64 | 0.57 | 0.65 |
| GSE207248 \| Pac + H + Pert \| pCR | 0.56 | 0.63 | 0.59 | 0.55 | | 0.60 |
| GSE123845 \| AC + taxane (+/−H) \| pCR | 0.54 | 0.58 | 0.63 | 0.60 | 0.71 | 0.62 |
| GSE41998 / Test 1 \| AC −> Pac \| pCR | 0.73 | 0.72 | 0.72 | 0.66 | 0.69 | 0.67 |
| GSE41998 / Test 2 \| AC −> Ixa \| pCR | 0.66 | 0.68 | 0.67 | 0.62 | 0.65 | 0.60 |
| GSE41998 / Test 3 \| AC phase \| CR/PR | 0.65 | 0.65 | 0.63 | | 0.57 | 0.58 |
| GSE32646 \| Pac −> FEC \| pCR | 0.79 | 0.77 | 0.76 | 0.76 | 0.70 | 0.71 |
| GSE20194 \| Pac + FAC \| pCR [E] | 0.81 | 0.81 | 0.81 | 0.75 | 0.77 | 0.74 |
| GSE23988 \| FAC −> Doc/Cape \| pCR [E] | 0.81 | 0.78 | 0.77 | 0.77 | 0.73 | 0.81 |
| GSE22226 \| AC + taxane (unspecified) \| pCR | 0.67 | 0.67 | 0.68 | | 0.70 | 0.67 |
| GSE25055 \| Taxane/anthracycline \| pCR [E] | 0.79 | 0.79 | 0.80 | 0.78 | 0.77 | 0.76 |
| GSE25066 \| Pac−containing subset \| pCR | 0.67 | 0.67 | 0.66 | 0.66 | | 0.67 |
| GSE20271 / Test 1 \| Pac + FAC \| pCR | 0.54 | 0.58 | 0.64 | 0.56 | 0.61 | 0.73 |
| GSE20271 / Test 2 \| FAC \| pCR | 0.44 | 0.59 | 0.63 | | 0.64 | 0.47 |
| GSE194040 / Test 1 \| Pac + Ganitumab −> AC / TNBC \| pCR | 0.34 | 0.41 | 0.46 | 0.47 | 0.44 | 0.54 |
| GSE194040 / Test 2 \| Pac + Ganitumab −> AC / HR+/HER2− \| pCR | 0.50 | 0.57 | 0.60 | 0.61 | 0.71 | 0.70 |
| GSE194040 / Test 3 \| Pac + Ganetespib −> AC / TNBC \| pCR | 0.61 | 0.62 | 0.63 | 0.65 | 0.71 | 0.56 |
| GSE194040 / Test 4 \| Pac + Ganetespib −> AC / HR+/HER2− \| pCR | 0.55 | 0.50 | 0.52 | 0.49 | 0.55 | 0.56 |
| GSE194040 / Test 5 \| Pac + MK−2206 −> AC / TNBC \| pCR | 0.57 | 0.58 | 0.65 | 0.73 | 0.70 | 0.51 |
| GSE194040 / Test 6 \| Pac + MK−2206 −> AC / HR+/HER2− \| pCR | 0.30 | 0.34 | 0.38 | 0.55 | 0.27 | 0.30 |
| GSE194040 / Test 7 \| Pac + Pembrolizumab −> AC / TNBC \| pCR | 0.69 | 0.64 | 0.66 | 0.54 | 0.53 | 0.41 |
| GSE194040 / Test 8 \| Pac + Pembrolizumab −> AC / HR+/HER2− \| pCR | 0.70 | 0.69 | 0.67 | 0.63 | 0.65 | 0.66 |
| GSE194040 / Test 9 \| Pac −> AC / TNBC \| pCR | 0.60 | 0.58 | 0.57 | 0.58 | 0.48 | 0.57 |
| GSE194040 / Test 10 \| Pac −> AC / HR+/HER2− \| pCR | 0.75 | 0.71 | 0.73 | 0.70 | 0.69 | 0.75 |
| GSE194040 / Test 11 \| Pac + AMG 386 −> AC / TNBC \| pCR | 0.51 | 0.50 | 0.51 | 0.49 | 0.48 | 0.45 |
| GSE194040 / Test 12 \| Pac + AMG 386 −> AC / HR+/HER2− \| pCR | 0.72 | 0.78 | 0.79 | 0.75 | 0.64 | 0.70 |
| GSE194040 / Test 13 \| Pac + MK−2206 + H −> AC / HER2+ \| pCR | 0.55 | 0.49 | 0.49 | 0.52 | 0.45 | 0.41 |
| GSE194040 / Test 14 \| Pac + Neratinib −> AC / TNBC \| pCR | 0.51 | 0.63 | 0.64 | 0.67 | 0.72 | 0.60 |
| GSE194040 / Test 15 \| Pac + Neratinib −> AC / HR+/HER2− \| pCR | 0.23 | 0.14 | 0.17 | 0.19 | 0.36 | 0.29 |
| GSE194040 / Test 16 \| Pac + Neratinib −> AC / HER2+ \| pCR | 0.54 | 0.58 | 0.61 | 0.59 | 0.60 | 0.59 |
| GSE194040 / Test 17 \| Pac + Veli + Carbo −> AC / TNBC \| pCR | 0.64 | 0.59 | 0.59 | 0.52 | 0.68 | 0.64 |
| GSE194040 / Test 18 \| Pac + Veli + Carbo −> AC / HR+/HER2− \| pCR | 0.91 | 0.94 | 0.96 | 0.93 | 0.94 | 0.92 |
| GSE194040 / Test 19 \| Pac + H −> AC / HER2+ \| pCR | 0.78 | 0.73 | 0.64 | 0.61 | 0.48 | 0.56 |
| GSE194040 / Test 20 \| T−DM1 + Pert −> AC / HER2+ \| pCR | 0.81 | 0.76 | 0.73 | | 0.61 | 0.64 |
| GSE194040 / Test 21 \| Pac + Pert + H −> AC / HER2+ \| pCR | 0.70 | 0.69 | 0.70 | 0.64 | 0.52 | 0.69 |
| GSE194040 / Test 22 \| Pac + AMG 386 + H −> AC / HER2+ \| pCR | 0.53 | 0.43 | 0.42 | 0.49 | 0.43 | 0.38 |

Patient−level, training−fold−fitted CV. Red = 0; white = 0.5; blue = 1. Grey = unavailable.
