## Supplementary Files Figures And Tables for "Multi-objective domain adaptation for pan-cancer transfer of pharmacogenomic drug response to patient tumours": Supplementary_File S1.docx

Supplementary File S1
Mathematical formulation and joint training of MODAE+

### Mathematical formulation of MODAE+

Let $x_{i}^{cl}\in\mathbb{R}^{p}$ denote the gene-expression profile of cell line $i$, and let $x_{j}^{pt}\in\mathbb{R}^{p}$ denote the gene-expression profile of patient tumour $j$. Let $r_{i}\in\mathbb{R}^{D}$ represent the drug-sensitivity vector for cell line $i$, where $D$ is the number of drugs. Patient survival time and event indicator are denoted by $T_{j}$ and $E_{j}$, respectively. Cancer-lineage labels for cell lines and patients are denoted by $c_{i}^{cl}$ and $c_{j}^{pt}$ when available. Cell-line cancer-lineage annotations were obtained from DepMap using OncoTree labels, whereas TCGA tumour types were manually mapped to the same ontology and harmonized at OncoTree Level 1.

All inputs are mapped to a latent representation through a shared encoder $E_{\theta}$:

$$z=E_{\theta}\left( x \right).$$

The latent representation is partitioned into shared and domain-specific components:

$$z=\left[ z^{sh},z^{pt},z^{cl} \right],$$

where $z^{sh}$ captures transferable biological information, while $z^{pt}$ and $z^{cl}$ represent patient-specific and cell-line-specific variation, respectively.

#### Masked reconstruction

Reconstruction is performed using domain-specific masked latent representations. For patient tumours and cell lines, the decoder input is defined as:

$$\tilde{z}^{pt}=\left[ z^{sh},z^{pt},0 \right], \tilde{z}^{cl}=\left[ z^{sh},0,z^{cl} \right].$$

The reconstructed expression profiles are obtained as:

$$\hat{x}^{pt}=D_{\theta}\left( \tilde{z}^{pt} \right), \hat{x}^{cl}=D_{\theta}\left( \tilde{z}^{cl} \right).$$

Reconstruction losses are computed separately for patient and cell-line samples using mean squared error normalized by input dimension:

$$L_{rec}^{pt}=\frac{1}{n_{pt}p}\sum_{j=1}^{n_{pt}} \parallel x_{j}^{pt}-\hat{x}_{j}^{pt}\parallel_{2}^{2}, L_{rec}^{cl}=\frac{1}{n_{cl}p}\sum_{i=1}^{n_{cl}} \parallel x_{i}^{cl}-\hat{x}_{i}^{cl}\parallel_{2}^{2}.$$

#### Adversarial domain alignment and deconfounding

Shared latent representations are aligned using a Wasserstein domain critic $C_{dom}$. The critic is trained to distinguish patient and cell-line shared embeddings:

$$L_{critic}=\frac{1}{n_{pt}}\sum_{j=1}^{n_{pt}} C_{dom}\left( z_{j}^{pt,sh} \right)-\frac{1}{n_{cl}}\sum_{i=1}^{n_{cl}} C_{dom}\left( z_{i}^{cl,sh} \right)+\lambda_{gp}L_{gp},$$

where $L_{gp}$ is the gradient-penalty term:

$$L_{gp}=\mathbb{E}_{\hat{z}}\left[ \left( \parallel\nabla_{\hat{z}}C_{dom}\left( \hat{z} \right)\parallel_{2}-1 \right)^{2} \right].$$

The encoder is trained adversarially to reduce domain-specific information in the shared latent representation. The adversarial alignment loss for the encoder is defined as:

$$L_{adv}=\frac{1}{n_{cl}}\sum_{i=1}^{n_{cl}} C_{dom}\left( z_{i}^{cl,sh} \right).$$

A deconfounding penalty is applied to encourage separation between shared and private components:

$$L_{decf}=\parallel\left( Z^{pt,sh} \right)^{T}Z^{pt,pt}\parallel_{F}^{2}+\parallel\left( Z^{cl,sh} \right)^{T}Z^{cl,cl}\parallel_{F}^{2}.$$

#### Drug-response prediction

Drug sensitivity is predicted from the shared cell-line latent representation using a multi-channel regression head:

$$\hat{r}_{i}=P_{drug}\left( z_{i}^{cl,sh} \right).$$

The drug-response loss is a masked mean squared error over observed drug–cell-line pairs:

$$L_{drug}=\frac{\sum_{i=1}^{n_{cl}} \sum_{d=1}^{D} m_{id}\left( r_{id}-\hat{r}_{id} \right)^{2}}{\sum_{i=1}^{n_{cl}} \sum_{d=1}^{D} m_{id}},$$

where $m_{id}=1$ if drug sensitivity is observed for cell line $i$ and drug $d$, and $m_{id}=0$ otherwise. The multi-channel architecture assigns drug-specific hidden units before the final output layer to reduce excessive correlation between predictions for different drugs.

#### Survival prediction

Patient survival is modelled using a Cox proportional hazards objective. A survival head maps the shared patient embedding to a risk score:

$$\eta_{j}=P_{surv}\left( z_{j}^{pt,sh} \right).$$

The survival loss is the negative Cox partial log-likelihood:

$$L_{surv}=-\sum_{j:E_{j}=1} \left[ \eta_{j}-log\left( \sum_{k\in R\left( T_{j} \right)} \exp\left( \eta_{k} \right) \right) \right],$$

where $R\left( T_{j} \right)$ denotes the risk set at time $T_{j}$. In implementation, ties were handled using the Efron approximation.

#### Cancer-lineage classification

MODAE+ incorporates supervised cancer-lineage classification to preserve biologically meaningful organization in the shared latent space. The classifier predicts cancer-lineage labels from the shared embeddings:

$$\hat{c}_{j}^{pt}=P_{lineage}\left( z_{j}^{pt,sh} \right), \hat{c}_{i}^{cl}=P_{lineage}\left( z_{i}^{cl,sh} \right).$$

Patient and cell-line lineage-classification losses are computed separately using masked multi-class cross-entropy:

$$L_{lineage}^{pt}=-\frac{1}{\left| I_{pt} \right|}\sum_{j\in I_{pt}} \sum_{k=1}^{K} \mathbb{I}\left( c_{j}^{pt}=k \right)\log\hat{c}_{jk}^{pt},$$

$$L_{lineage}^{cl}=-\frac{1}{\left| I_{cl} \right|}\sum_{i\in I_{cl}} \sum_{k=1}^{K} \mathbb{I}\left( c_{i}^{cl}=k \right)\log\hat{c}_{ik}^{cl},$$

where $I_{pt}$ and $I_{cl}$ denote labelled patient and cell-line samples, respectively. This objective encourages samples belonging to the same cancer lineage to cluster together in the shared latent space while preserving separation between biologically distinct lineages.

#### Multi-objective FAMO optimization

The complete MODAE+ objective consists of eight loss components:

$$L_{t}=\left[ L_{rec}^{pt},L_{rec}^{cl},L_{adv},L_{decf},L_{surv},L_{drug},L_{lineage}^{pt},L_{lineage}^{cl} \right].$$

These objectives are optimized using Fast Adaptive Multitask Optimization (FAMO), which dynamically adjusts the contribution of each objective during training. Adaptive objective weights are obtained from trainable parameters $\rho_{t}$:

$$w_{t}=softmax\left( \rho_{t} \right).$$

The FAMO-weighted gradient used to update the MODAE+ model parameters is:

$$G_{\theta}=\sum_{k=1}^{8} \frac{c_{t}w_{k,t}}{L_{k,t}}\nabla_{\theta}L_{k,t}, c_{t}=\left( \sum_{k=1}^{8} \frac{w_{k,t}}{L_{k,t}} \right)^{-1}.$$

After updating the model parameters, the FAMO parameters are optimized according to the change in logarithmic loss values:

$$\Delta\log L_{t}=logL_{t}-logL_{t+1}.$$

The weight parameters are updated as:

$$\rho_{t+1}\leftarrow\mathrm{Adam}_{weights}\left( \rho_{t},J_{w}^{T}\Delta\log L_{t},\gamma\right),$$

where $J_{w}$ is the Jacobian of the normalized weights with respect to $\rho$, and $\gamma$ denotes the weight-decay parameter. This adaptive optimization strategy dynamically balances competing objectives during training, preventing individual losses from dominating the optimization process while improving convergence compared with fixed-weight formulations.

### Joint training algorithm

**Algorithm 1. Joint training of MODAE+**

**Require:** Patient expression $X_{pt}$; cell-line expression $X_{cl}$; cell-line drug sensitivity $R_{cl}$; patient survival times $T_{pt}$ and event indicators $E_{pt}$; patient and cell-line lineage labels $C_{pt}$ and $C_{cl}$; epochs $N$; model mini-batches per epoch $M$; adversarial mini-batches per model mini-batch $m$.

**Require:** Encoder $E$; decoder $D$; critic $C_{dom}$; multi-channel drug predictor $P_{drug}$; survival predictor $P_{surv}$; lineage classifier $P_{lineage}$; model, adversarial and FAMO weight optimizers.

1. Initialize $\theta=\{E,D,P_{drug},P_{surv},P_{lineage}\}$, critic parameters $\theta_{dom}$ and FAMO parameters $\rho$.
2. Initialize $w=softmax\left( \rho\right)$.
3. **for** epoch $=1,\ldots,N$ **do**
4. **for** $b=1,\ldots,M$ **do**
5. **Adversarial domain-critic update**
6. **for** $b_{a}=1,\ldots,m$ **do**
7. Sample mini-batches $X_{pt}^{b_{a}}$ and $X_{cl}^{b_{a}}$.
8. Encode $Z_{pt}=E\left( X_{pt}^{b_{a}} \right)$ and $Z_{cl}=E\left( X_{cl}^{b_{a}} \right)$.
9. Extract $Z_{pt}^{sh}=f_{sh}\left( Z_{pt} \right)$ and $Z_{cl}^{sh}=f_{sh}\left( Z_{cl} \right)$.
10. Compute the Wasserstein critic loss with gradient penalty:

$$L_{critic}=\frac{1}{\left| pt \right|}\sum_{pt} C_{dom}\left( Z_{pt}^{sh} \right)-\frac{1}{\left| cl \right|}\sum_{cl} C_{dom}\left( Z_{cl}^{sh} \right)+\lambda_{gp}L_{gp}.$$

11. Update $\theta_{dom}^{t+1}\leftarrow\mathrm{Adam}_{adv}\left( \theta_{dom}^{t},\nabla_{\theta_{dom}}L_{critic} \right)$.
12. **end for**
13. **MODAE+ model update**
14. Sample matched mini-batches $\left( X_{pt}^{b},T_{pt}^{b},E_{pt}^{b},C_{pt}^{b} \right)$ and $\left( X_{cl}^{b},R_{cl}^{b},C_{cl}^{b} \right)$.
15. Encode $Z_{pt}=E\left( X_{pt}^{b} \right)$ and $Z_{cl}=E\left( X_{cl}^{b} \right)$.
16. Split into shared and private components:

$$Z_{pt}=\left[ Z_{pt}^{sh},Z_{pt}^{pt},Z_{pt}^{cl} \right], Z_{cl}=\left[ Z_{cl}^{sh},Z_{cl}^{pt},Z_{cl}^{cl} \right].$$

17. Apply domain-specific masking:

$$\tilde{Z}_{pt}=\left[ Z_{pt}^{sh},Z_{pt}^{pt},0 \right], \tilde{Z}_{cl}=\left[ Z_{cl}^{sh},0,Z_{cl}^{cl} \right].$$

18. Reconstruct $\hat{X}_{pt}=D\left( \tilde{Z}_{pt} \right)$ and $\hat{X}_{cl}=D\left( \tilde{Z}_{cl} \right)$.
19. Predict drug sensitivity, survival risk and cancer lineage:

$$\hat{R}_{cl}=P_{drug}\left( Z_{cl}^{sh} \right), \hat{\eta}_{pt}=P_{surv}\left( Z_{pt}^{sh} \right),$$

$$\hat{C}_{pt}=P_{lineage}\left( Z_{pt}^{sh} \right), \hat{C}_{cl}=P_{lineage}\left( Z_{cl}^{sh} \right).$$

20. Compute the eight-objective vector:

$$L_{t}=\left[ L_{rec}^{pt},L_{rec}^{cl},L_{adv},L_{decf},L_{surv},L_{drug},L_{lineage}^{pt},L_{lineage}^{cl} \right].$$

- Here, the components are patient and cell-line reconstruction, adversarial alignment, deconfounding, Cox survival, masked drug sensitivity, and patient and cell-line masked lineage classification.

21. Compute $w_{t}=softmax\left( \rho_{t} \right)$.
22. Compute the FAMO-weighted model gradient:

$$G_{\theta}=\sum_{k=1}^{8} \frac{c_{t}w_{k,t}}{L_{k,t}}\nabla_{\theta}L_{k,t}, c_{t}=\left( \sum_{k=1}^{8} \frac{w_{k,t}}{L_{k,t}} \right)^{-1}.$$

23. Update $\theta_{t+1}\leftarrow\mathrm{Adam}_{joint}\left( \theta_{t},G_{\theta} \right)$.
24. Compute $\Delta\log L_{t}=logL_{t}-logL_{t+1}$.
25. Update $\rho_{t+1}\leftarrow\mathrm{Adam}_{weights}\left( \rho_{t},J_{w}^{T}\Delta\log L_{t},\gamma\right)$.
26. **end for**
27. **end for**

**Ensure:** Optimized parameters $\theta$ and $\theta_{dom}$; shared embeddings $Z^{sh}$; transferred patient drug-sensitivity estimates $\hat{R}_{pt}=P_{drug}\left( Z_{pt}^{sh} \right)$; survival risk predictions $P_{surv}\left( Z_{pt}^{sh} \right)$; and lineage predictions $P_{lineage}\left( Z^{sh} \right)$.
