## Supplementary figures and images for "Multi-objective domain adaptation for pan-cancer transfer of pharmacogenomic drug response to patient tumours"

### Supplementary Figure S1.png

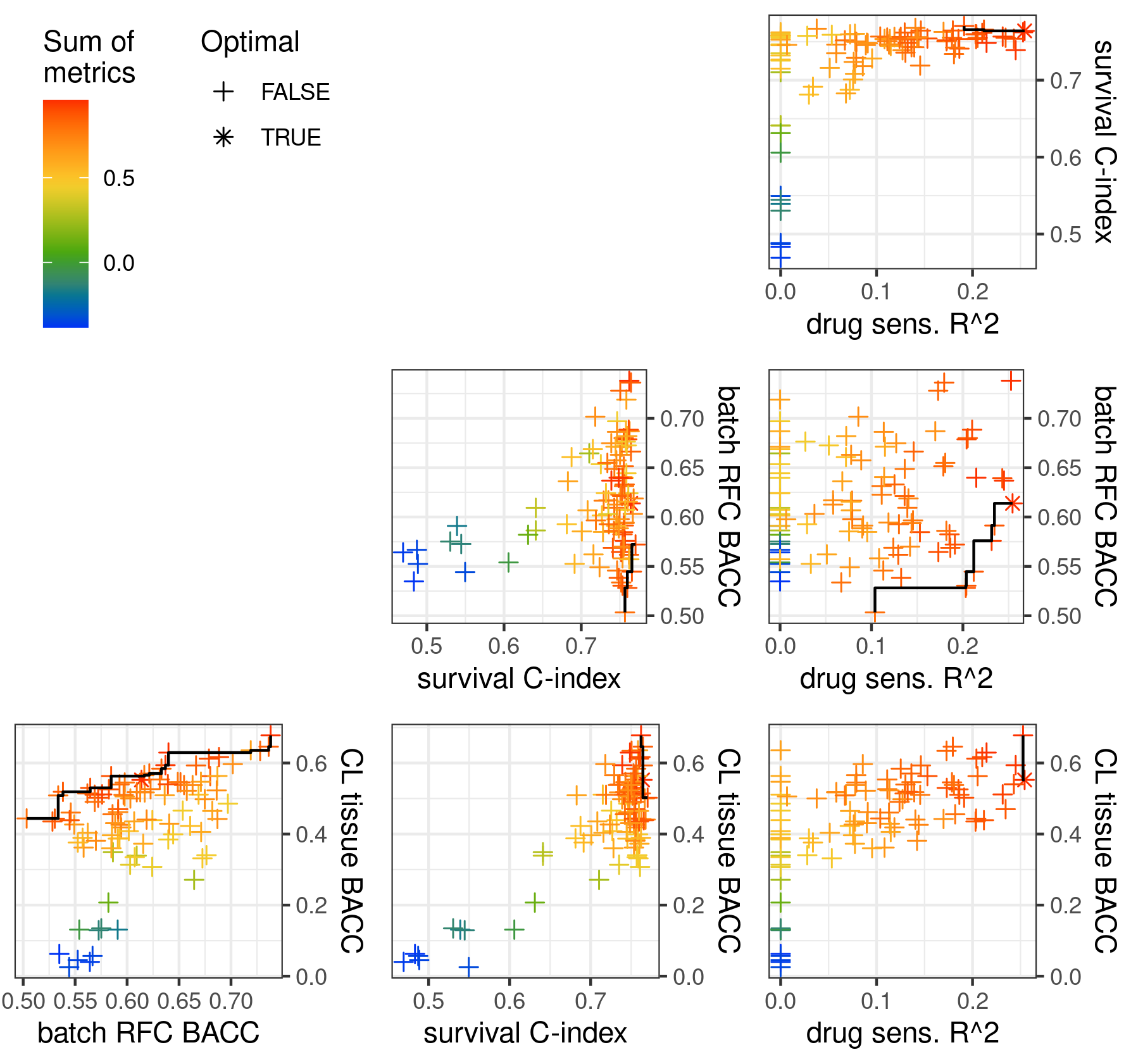
